# CD8 T cells with classical and NK-like cytotoxic gene expression programs mediate control of HBV replication and functional cure

**DOI:** 10.64898/2026.06.27.732784

**Authors:** Elias Isaac Lattouf, Martin Feuerherd, Lea M Bartsch, Hannah K Drescher, Suzan Dijkstra, Martin A Villanueva, Ruben C Hoogeveen, David Lieb, Koen Van Den Berge, Ewoud De Troyer, Sonu Subudhi, Alex S Genshaft, Nádia Conceição-Neto, Anna K Traunbauer, Aljawharah Alrubayyi, Charles R Crain, Loghman Salimzadeh, Marcos Damasio, Boris J Beudeker, Danie PM La, Juan Diego Sanchez Vasquez, James A Cheney, Mantero V Moreno-Cheek, Stuti Shah, Jasneet Aneja, Michael T Waring, Nadia Alatrakchi, Arthur Y Kim, Robert J de Knegt, Lia L Lewis-Ximenez, Jeroen Aerssens, Jacques Bollekens, Nir Hacohen, Gaurav D Gaiha, Raymond T Chung, Jordan J Feld, Harry LA Janssen, Alex K Shalek, Andre Boonstra, Adam J Gehring, Georg M Lauer

## Abstract

Chronic hepatitis B is characterized by a decades-long evolving engagement between host immunity and the hepatitis B virus (HBV). Understanding the molecular characteristics of HBV-specific CD8 T cells linked to control of viral replication and antigenemia is essential to design effective immunotherapeutic modalities. Here we show that HBV-specific CD8 T cells, even during infection stages with extremely high viral loads, lack the features of terminally exhausted CD8 T cells observed in chronic HCV and HIV infection or cancer. Instead, we observe emerging gene expression programs over disease stages that correlate with increasing HBV control, which include a *bona fide* cytotoxic and T-cell localization program associated with low levels of viral replication, and a second *NK-like* T-cell program that combines expression of classical NK markers (*KIR*s, *KLR*s, *FCGR3A*, *TYROBP*, *IKZF2*) with cytotoxic genes (*GZMB*, *GNLY*, *PRF1*), which emerges with complete control of HBV viremia and antigenemia. We also found enrichment of both CD8 T-cell programs in HIV-specific CD8 T cells from HIV elite controllers, supporting a conserved role in controlling persistent viral infections with viral reservoirs.

## Introduction

Chronic hepatitis B (CHB) due to infection with hepatitis B virus (HBV) continues to be a major clinical challenge worldwide, affecting 250 million people and resulting in 800,000 deaths annually^1^. While antiviral therapy of CHB effectively suppresses viral replication and transmission, carries a favorable side-effect profile, and significantly reduces morbidity and mortality, treatment is lifelong in most cases as viremia recurs after cessation of therapy, often leading to liver disease^2^. Delivering uninterrupted treatment remains difficult in many regions with the highest prevalence: moreover, infection and ongoing care may lead to social stigma. Thus, a short-term finite treatment that achieves complete HBV clearance would have a major impact on the global burden of CHB. Immunological containment with fully suppressed viral replication and antigenemia is the most promising path to sustained HBV cure. During natural infection spontaneous and durable control is the outcome of almost all adult-acquired HBV infections, and even occurs in about 1% of people with infant-acquired infections each year after decades of chronic infection, with or without treatment^3^.

Importantly, HBV DNA persists in the form of covalently closed circular DNA (cccDNA) and integrated DNA in hepatocytes, even when assays for both HBV DNA and Hepatitis B s antigen (HBsAg) in the blood are negative^4^. This is distinct from hepatitis C virus (HCV) infection, which lacks a DNA intermediate and can be completely eliminated both immunologically and through antiviral treatment^5^. Natural viral containment of HBV is more similar to the situation of elite controllers in human immunodeficiency virus (HIV) infection, who have undetectable HIV viremia in the blood and do not progress to immunodeficiency in the absence of antiviral treatment despite an ongoing viral reservoir^6^. Most novel interventions for CHB therefore aim to induce such a state of immunological control, but very few studies have fully dissected host immune responses in this scenario, whether in blood or liver. Consequently, the specific immunological correlates that distinguish CHB functional cure (FC) – with stably negative HBsAg and HBV viral load (VL) off treatment – remain poorly defined^7,8^.

Another highly unusual feature of CHB, distinguishing it from other chronic viral infections, is that most people living with hepatitis B develop immunological control of HBV decades into infection, transitioning from viral loads greater than 10^8^ to less than 2,000 IU/mL, although only a small subgroup will eventually achieve sustained loss of detectable serum HBsAg^9,10^. This is very different from untreated HCV or HIV, where early viral setpoints remain mostly unchanged or increase over decades^11,12^, or from cytomegalovirus (CMV) or Epstein-Barr virus (EBV) where viremia is contained shortly after infection^13,14^. This suggests evolution of HBV immunity over time towards more effective states. Furthermore, as the reduction in viral replication is not necessarily paralleled by a similar reduction in circulating HBsAg, it is likely that the correlates of control of HBV replication and HBsAg are not identical^2^.

HBV-specific CD8 T-cell responses are thought to be essential for controlling HBV replication as well as antigenemia^15^, but the particular characteristics of protective HBV-specific CD8 immunity as well as the molecular mechanisms subverting its efficacy across different stages of CHB await full definition. To address this knowledge gap, a multinational consortium designed a multicenter study collecting liver and blood samples across all stages of CHB and FC. These stages, including FC are observed over the course of a lifetime of infection, and their inclusion enables a precise and deep characterization of associated HBV-specific CD8 T-cell immunity. We also assessed HBV-specific CD8 T cells from individuals with adult-acquired acute hepatitis B who naturally control HBV within 6 months for comparison. We identify two complex cytotoxic gene expression programs (GEPs) in HBV-specific CD8 T cells: one that increases with better control of viral replication, and a signature of cytotoxic T cells with NK-like properties that arises in HBV-specific cells after complete control of HBV replication and antigenemia. Moreover, we observe overlap of GEPs enriched in HIV-specific CD8 T cells of HIV elite controllers, suggesting a broader relevance of virus-specific CD8 T cells with these distinct cytotoxic GEPs for control of persistent viral infections and as novel targets for immune-based antiviral interventions.

## Results

### Cohorts representing all clinical scenarios of HBV infection

Different clinical phases of CHB are characterized by vastly distinct levels of viral replication together with particular patterns of HBV antigenemia and liver inflammation. Our consortium collected peripheral blood mononuclear cells (PBMC) and intrahepatic lymphocytes (IHL) from liver fine needle aspirates (FNA) in a cross-sectional cohort of 68 untreated individuals with CHB and 27 participants who naturally achieved FC (Extended Data Fig. 1a). The cohort spans all clinical stages of CHB, with viral loads ranging from > 10^8^ IU/mL to undetectable and a broad range of HBsAg levels, enabling analysis of CD8 T-cell immunity over the full range of viral replication and control. As the clinical evolution of CHB through these phases typically unfolds over decades and only a small fraction of people living with hepatitis B eventually achieves complete control of viral replication and antigenemia, we were unable to study individuals longitudinally throughout their infection course. As such, the cross-sectional cohort captures the known phases of CHB^2,8^ and, consistent with the natural history of predominantly perinatally acquired CHB, median age increased from 29 years in participants with Hepatitis B e antigen (HBeAg)-positive infection, to 47 years in HBeAg-negative infection, to 60 years in FC (Extended Data Fig. 1b). We complemented this cross-sectional cohort of untreated CHB and FC, with longitudinal samples collected before and after interruption of long-term CHB antiviral treatment as well as from individuals followed throughout acute adult-acquired HBV infection. Collectively, these cohorts encompass the full spectrum of clinically observed HBV infection states and outcomes (Extended Data Fig. 1c).

Based on the expression of HLA alleles matching an extensive library of HBV HLA class I multimers (Supplementary Table 1), we were able to screen PBMC of 84 individuals for the presence of HBV-specific CD8 T-cell responses (Extended Data Fig. 2a). Samples with sufficiently large HBV-specific CD8 T-cell populations were index-sorted (Extended Data Fig. 2b) directly *ex vivo* followed by scRNA-seq using the Smart-seq2 protocol^16,17^ (Extended Data Fig. 2c), enabling analysis of data from 39 untreated CHB+FC participants (Fig. 1a,b, Extended Data Fig. 2d; see Supplementary Table 2).

**Fig 1.**
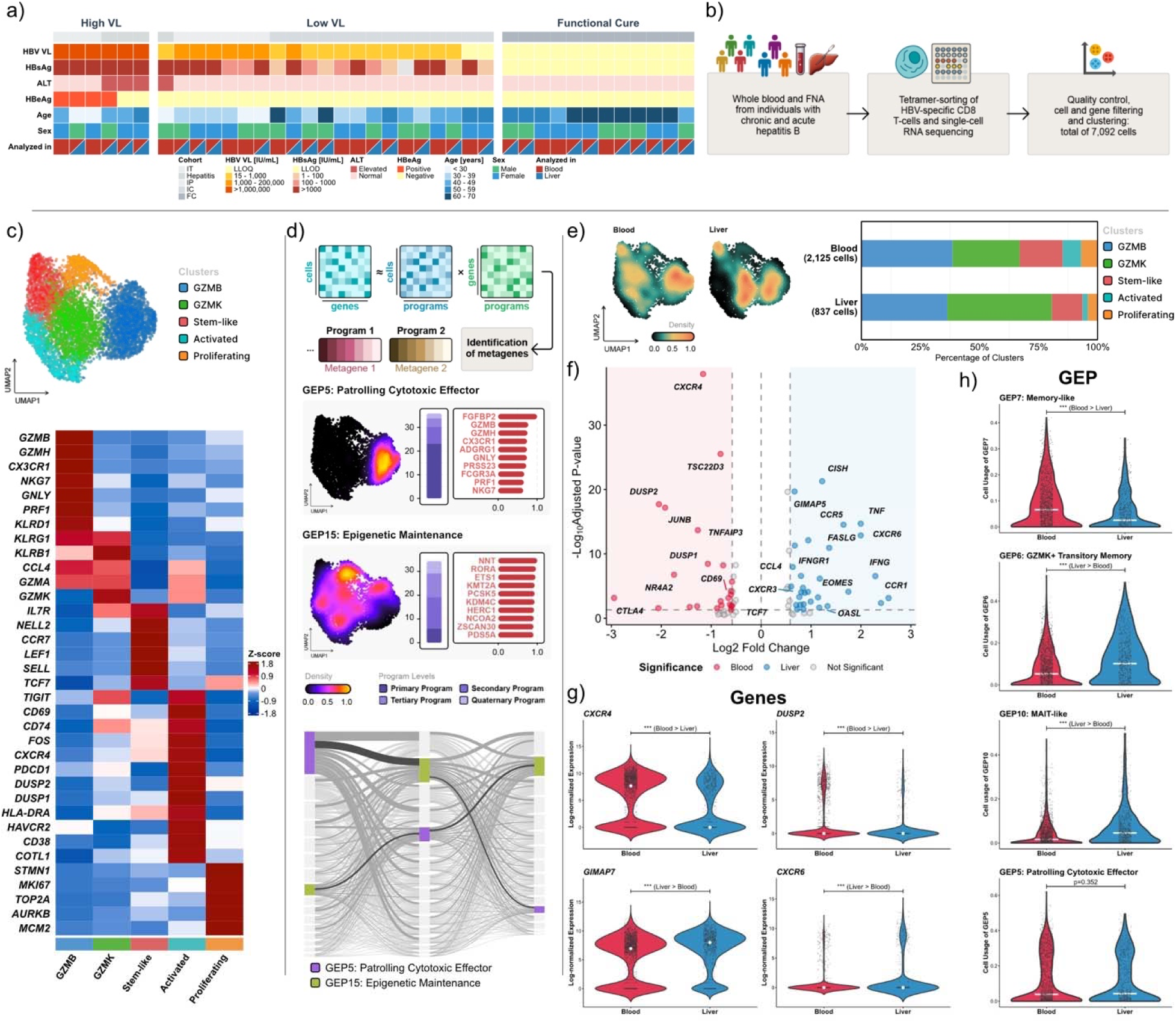
Diversity of peripheral and intrahepatic HBV-specific CD8 T cells over the whole breadth of CHB. (**a**) Clinical values of participants (n = 39) with CHB and high viral load, low viral load and FC. (**b**) Overview of the workflow to assess the participants’ HBV-specific T-cell repertoire. (**c**) UMAP-based Leiden-clustering with 5 clusters of prototypic T-cell immune statuses across the whole breadth of HBV infection (7,092 cells, n = 52 participants) defined by their DEG as shown in the heatmap. Gene expression is depicted as z-score. (**d**) cNMF identifies gene expression programs (GEP) based on ‘cell x gene’-matrix with each cell having a usage value of each program (‘cell x GEP’-matrix) and each program having a weight of each gene (‘gene x GEP’-matrix) in the dataset. UMAP shows density of cells with high usage of indicated example GEPs (*Patrolling Cytotoxic Effector*, and *Epigenetic Maintenance*). Hierarchical stacked bar chart shows abundance of cells with the respective program as their primary highest usage, secondary, tertiary, or quaternary GEP. Horizontal bar graph shows top genes and gene weights defining the respective GEP. Sankey plot indicates the interconnectivity of each program with each other programs based on each singular cell’s hierarchy as primary, secondary, and tertiary programs for each cell. (**e**) Density of HBV-specific CD8 T cells from blood and liver are depicted on a UMAP. The distribution of cells from participants with paired blood and liver is shown over Leiden clusters (2,962 cells, n = 27 participants). (**f**) Volcano plot depicts DGE between cells from participants with paired blood (red) and liver (blue) cells. Genes of interest with > 1.5-fold change and adjusted *p*-value < 0.05 are labeled. (**g**) Violin plots show expression of selected genes in HBV-specific CD8 T cells from blood and liver. (**h**) Violin plots show selected GEP usage between cells from blood and liver. Statistical significance was determined using Wilcoxon rank-sum test (unpaired, nonparametric, two-sided) and for DGE supplemented with Benjamini-Hochberg correction for multiple testing; \**p* ≤ 0.05, ** *p* ≤ 0.01, *** *p* ≤ 0.001, non-significant *p*-value is given if *p* > 0.05.

### Transcriptional profiles of HBV-specific CD8 T cells

Based on the single-cell transcriptomes of HBV-specific CD8 T cells from all PBMC and IHL samples (6,020 from blood and 1,072 from liver), Leiden clustering identified 5 distinct CD8 T-cell populations (Fig. 1c). Based on differential gene expression (DGE) we assigned 5 canonical T-cell differentiation states: stem-like, cytotoxic GZMB, GZMK, proliferating, and activated T cells (Fig. 1c)^18–20^. Previously, we and others described differences between HBV antigen-specificities in regards to their phenotype and function^21–24^, based on analyses of the two most studied HLA-A*0201 restricted CD8 T-cell responses: core18 (FLPSDFFPSV) and pol455 (GLSRYVARL). Here, we had the opportunity to analyze responses to 8 core epitopes (restricted by 6 alleles) and 5 pol epitopes (restricted by 4 alleles). In this broad analysis we confirm that core specificities express more *PDCD1* (encoding PD-1) as reported in previous studies on the two A*0201 responses^21,22^ as well as in a study comparing 1 core and 3 pol responses restricted by A*1101^25^. Beyond that, in contrast to the classic A*0201 responses^24^, core and pol-responses were equally distributed across the 5 clusters, suggesting a less universal dichotomy between core and pol responses targeting distinct epitopes restricted by various HLA alleles (Extended Data Fig. 3).

To achieve a more nuanced understanding of T-cell differentiation states not readily discerned by traditional clustering analysis approaches, we incorporated consensus non-negative matrix factorization (cNMF) to our analytic approaches^26^. cNMF goes beyond assessing individual expressed genes by identifying networks of co-expressed genes that represent gene expression programs (GEP) that define cell-lineage identities as well as cell activities, such as effector function or proliferation. Importantly, in contrast to traditional clustering, where each cell is only assigned to one cluster, cNMF can reveal a cell’s layered participation in multiple cellular states, as each cell can have multiple active GEPs. In our dataset, we defined 16 GEPs (Extended Data Fig. 4, Supplementary Table 3) corresponding to specific functional states such as cytotoxic effectors or central memory, but also GEPs indicative of more basic cellular processes such as transcriptional regulation or epigenetic maintenance that are operative in cells across several different clusters (Fig. 1d). As every cell can be part of multiple GEPs, we can define hierarchical layers of distinct cellular programs (Fig. 1d) for each individual cell, thereby creating a more complex understanding of antigen-specific CD8 T-cell regulation.

### Distinct and shared characteristics of HBV-specific CD8 T cells from the blood and the liver

Having access to both liver FNA and blood allowed us to directly contrast antigen-specific cells from the site of infection with those circulating in the blood in a paired comparison (Fig. 1e). That the HBV-specific CD8 T cells obtained from liver FNA were truly tissue-derived and not contaminating blood is supported by the much higher relative frequency of multimer-positive cells as a sign of compartmentalization of virus-specific CD8 T cells to the site of infection: we captured less than 3x HBV-specific CD8 T cells from the liver than from the blood despite having at least 2 magnitudes more input cells from the blood. This is in addition to their expression of the liver homing marker *CXCR6* and the homeostasis factor *GIMAP7* compared to cells derived from the blood that expressed higher levels of *CXCR4* and *TCF7* (Fig. 1f,g, Supplementary Table 4). Cells from FNA also showed much higher activity of the *MAIT-like* GEP (GEP10-*RORC, CXCR6, KLRB1*)^27,28^ as further evidence that FNA samples truly captured tissue-resident T cells and not contaminating blood (Fig. 1h). Apart from this typical liver-resident T-cell GEP, HBV-specific CD8 T cells from FNAs also displayed significantly increased activity of several other GEPs, including *GZMK+ Transitory Memory* (GEP6-*GZMK*, *CCL4*, *EOMES*) (Fig. 1h, Supplementary Fig. 1). By contrast, the *Memory-like* GEP (GEP7-*IL7R*, *CXCR4*, *TCF7*) was among those more utilized in T cells from the blood (Fig. 1h, Supplementary Fig. 1). While some GEPs like *Patrolling Cytotoxic Effector* (GEP5-*GZMB*, *GNLY*, *PRF1*) were not significantly enriched in either tissue, no GEP was exclusive to either liver- or blood-derived CD8 T cells, which might be explained by the unique open architecture of the liver vasculature and the large volume of blood flowing through it, allowing continuous exchange of cells between the compartments. Together, these data demonstrate significant differences in GEP dominance between HBV-specific CD8 T cells in liver versus blood.

### Both terminally differentiated and memory CD8 T cells are minor populations across all stages of CHB

Given the vastly different levels of HBV control across different stages of CHB, with viral loads ranging across 8 orders of magnitude, one would expect characteristic T-cell differentiation states at least in the most polarized viral load settings. Therefore, we analyzed the differentiation profiles of HBV-specific CD8 T cells across all CHB infection states but also in acute HBV and FC. HBV-specific CD8 T cells in treated and untreated CHB PD-1 and other immune checkpoints at varying levels and leading to the assumption that T-cell exhaustion is a key feature of CHB T-cell dysfunction^29^.

To delineate the differentiation of HBV-specific CD8 T cells, we first assessed the protein expression of CD127 and PD-1 (Fig. 2a) measured in our fluorescence index-sorting panel. The IL-7 receptor alpha-chain (CD127) demarcates both *bona fide* memory CD8+ T cells and the memory-like progenitor subset of exhausted T cells (Tpex), the latter of which sustains the long-term T-cell response and provides the proliferative burst following immune checkpoint inhibition^30^. While PD-1 is transiently upregulated upon TCR engagement during acute infection^31–33^, its sustained expression is a key feature of the distinct transcriptional (and epigenetic) program driven by chronic antigen exposure^34–36^. The biology of the phenotype and development of T-cell exhaustion – and the therapeutic potential of reinvigorating these cells with aPD1 blockade – was originally defined in the prototypic LCMV clone 13 mouse model of chronic viral infection^31,37^ and subsequently translated to reverse T-cell exhaustion in tumor-infiltrating lymphocytes in human cancer^38,39^. In humans, the natural history of HCV infection provides a paradigm for this dichotomy contrasting *bona fide* memory T cells following spontaneously resolving HCV with the establishment of terminal exhaustion during chronic HCV infection^36,40^.

**Fig 2.**
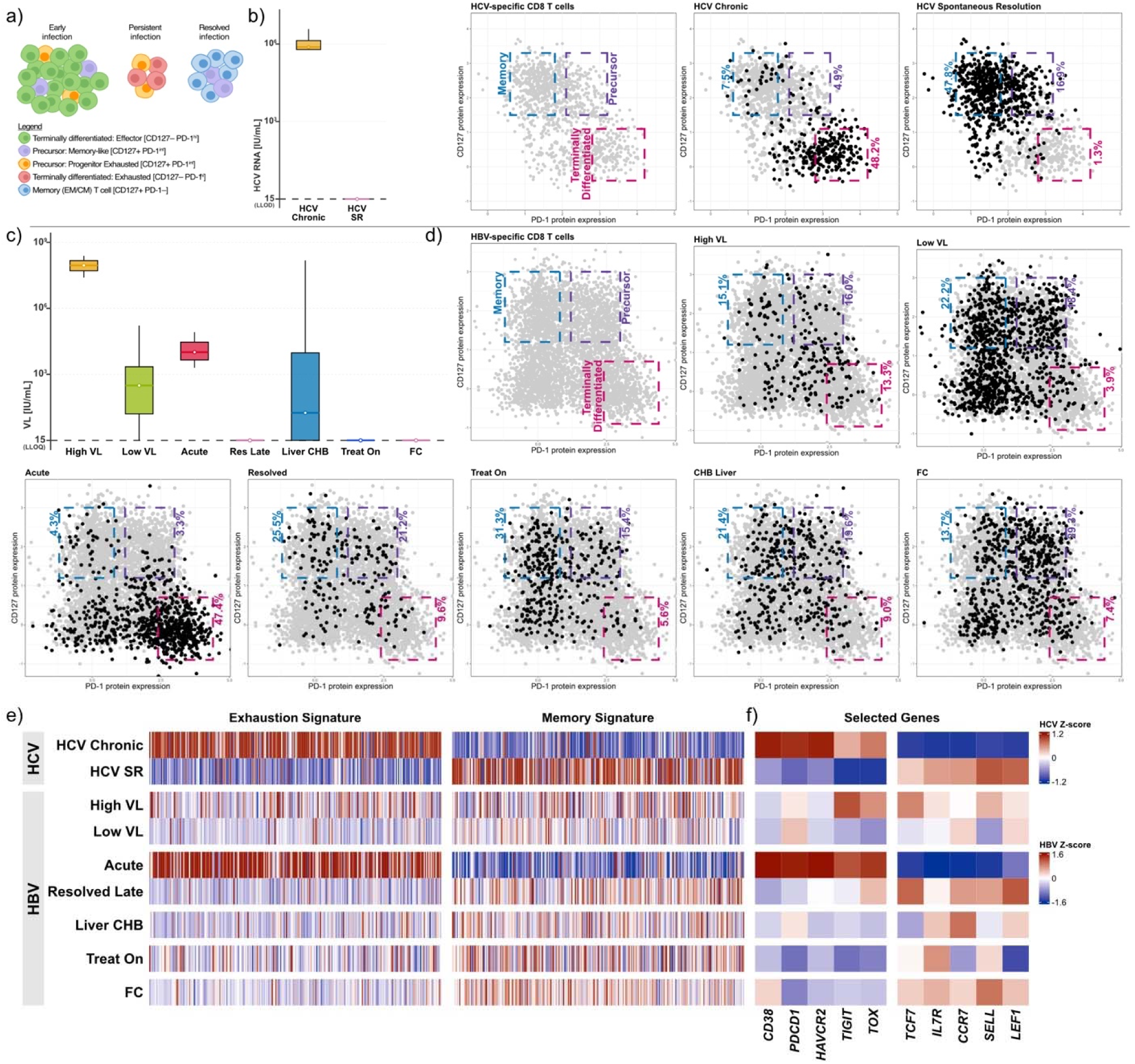
Evaluation of T-cell exhaustion in HCV- versus HBV-specific CD8 T cells. (**a**) Cartoon displays T-cell populations in Early infection, Persistent infection, and Resolved infection and indicates their CD127/PD-1 protein levels. (**b**) Boxplots indicate viremia of participants with chronic HCV (cells targeting conserved epitopes, n = 8) and with spontaneously resolved (SR) HCV infection (≥24 weeks post-resolution, n = 11). Flow panels show CD127 and PD-1 protein data of HCV-specific CD8 T cells (gray) with gates representing Memory (CD127+ PD-1-; blue), Precursors (CD127+ PD-1 mid; purple), and Terminally differentiated (CD127- PD-1hi; pink) T-cell populations. Protein data of HCV-specific CD8 T cells in the indicated cohort (black) versus all HCV-specific CD8 T cells (gray) are shown. (**c**) Boxplots indicate viremia of participants with hepatitis B (participants with CHB with High VL (n = 6), Low VL (n = 20), Acute HBV infection (n = 8), Acute Resolved (n = 8), CHB who contributed liver FNA (n = 19), CHB on treatment (Treat On, n = 5), and FC (n = 12)). (**d**) Flow panels show CD127 and PD-1 protein data of HBV-specific CD8 T cells (gray) with gates representing Memory (CD127+ PD-1-; blue), Precursors (CD127+ PD-1 mid; purple), and Terminally differentiated (CD127- PD-1hi; pink) T-cell populations. Protein data of HBV-specific CD8 T cells in the indicated cohort (black) versus all HBV-specific CD8 T cells (gray) are shown. (**e**) Heatmap showing the gene expression signatures of *bona fide* exhausted versus memory T cells (Top250 genes of bulk transcriptomes previously published by Tonnerre *et al.*^36^) between HCV-specific and HBV-specific CD8 T cells in the indicated cohorts. Gene expression is indicated as z-score. (**f**) Heatmap with gene expression of canonical exhaustion and memory markers in HCV-specific and HBV-specific CD8 T cells. Gene expression is indicated as z-score.

Therefore, we compared the protein levels of PD-1/CD127 in HBV-specific CD8 T cells with HCV-specific CD8 T cells in chronic and spontaneously resolved HCV infection (Fig. 2b). Surprisingly, we found very few terminally differentiated T-cells (PD-1hi/CD127-) in CHB, with a dominance of this phenotype only during early acute HBV infection (Fig 2c,d). By contrast, chronic HCV infection characterized by a major population of PD-1hi CD127- cells during persisting high viremia, despite CHB having decades of similarly high persisting viremia.

HBV-specific CD8 T cells in CHB had overall lower expression of PD-1 with or without concurrent expression of CD127 (Fig. 2d). As shown for CHB with low VL before^21^, all stages of CHB independent of the VL had a stable population of HBV-specific memory-like (PD-1+ CD127+) T cells that was comparable to a population found after spontaneous resolution of HCV infection (Fig. 2b-d). Nevertheless, a major population of HBV-specific CD8 T cells remained CD127 negative even at low or absent HBV viremia (CHB Low VL, and FC), in alignment with the fact that HBV, in contrast to HCV, is never completely eliminated^2,5^. Especially double negative antigen-specific T cells, a population almost non-existent in HCV, were enriched in controlled viremia and had a TEMRA phenotype throughout CHB and FC (Extended Data Fig. 5a,b). Overall, HBV-specific CD8 T-cell responses in blood and liver from chronic HBV show more complex and mostly PD-1 negative differentiation states compared to chronic HCV.

In order to define differentiation states on a broad molecular level, we compared the gene expression of HBV-specific CD8 T cells with previously published RNA-seq signatures of human T-cell exhaustion and *bona fide* memory signatures^36^. As expected, we found that the terminal T-cell exhaustion signature was enriched in HCV-specific CD8 T cells (to conserved epitopes) from chronic HCV infection, whereas cells from spontaneous resolution of acute HCV depicted high expression of the memory signature (Fig. 2e). Similar to the protein data, we did not observe enrichment of the terminally exhausted T-cell signature or memory profile in HBV-specific CD8 T cells across CHB and controlled HBV infection. Again, only T cells from early acute HBV infection expressed most genes from the exhaustion signature (Fig. 2e), which is explained by the overlap between this signature with highly activated, terminally differentiated effector T cells^41^. Surprisingly, blood-derived cells from CHB with high VL were lacking a clear exhaustion signature; while they expressed some characteristic genes such as *TOX* they also had high levels of *TCF7* (Fig. 2f). Even at the site of infection, HBV-specific CD8 T cells in CHB lacked the signature of terminally differentiated exhausted T cells. Importantly, the cells in the liver also did not express *TOX*, suggesting they were similar to tissue-resident memory cells instead of tissue-resident precursors to exhausted T cells which is the main population responding to PD-1 checkpoint inhibitors (Fig. 2e,f)^42^. Together, the protein and transcriptional data reveal that HBV-specific CD8 T cells across heterogeneous CHB states mostly occupy a surprisingly narrow corridor within the classical range of T-cell differentiation – in stark contrast to HCV infection – despite a comparable range of viremia across the HBV infection spectrum. The paucity of terminally differentiated and memory-like CD8 T cells expressing key exhaustion molecules in CHB is surprising and might explain why HBV viral replication is increasingly better controlled over decades^9,10^, but also why therapeutic responses to checkpoint inhibition are rare^43^.

### Differences in CD8 T-cell effector programs between CHB with high versus low viral replication

Based on the clinical observation that CHB evolves over decades from extremely high viral loads greater than 10^8^ to under 2,000 IU/mL in most individuals^9,10^, transcriptional differences of HBV-specific CD8 T cells were likely, despite the modest differences in basic differentiation states we detected above. Therefore, we compared the transcriptional landscapes of HBV-specific CD8 T cells from HBsAg-positive CHB individuals with high versus low HBV viral load (high ≥10^6^ IU/mL, median 7.7log_10_ versus low <10^6^ IU/mL, median 2.5log_10_) (Fig. 3a). Blood-derived HBV-specific CD8 T cells from participants in these two HBV viral load groups were enriched in different areas of the scRNA-seq Uniform Manifold Approximation and Projection (UMAP) (Supplementary Fig. 2) and differential gene expression (DGE) analysis identified 23 and 97 genes with significant upregulation between low and high levels of viral replication, respectively (Fig. 3b, Supplementary Table 4). Upregulated genes in low viral load included *CX3CR1*, *KLRB1*, *ZNF683*, *GZMB*, *GZMH*, *GNLY*, *NKG7* (Fig. 3b,c), typically observed in cytotoxic effector CD8 T cells. CD8 T cells in a high-viral-load environment expressed higher levels of T-cell activation markers including *CD74*, *HLA-DRA*, *RGS2*, *DUSP1*, *DUSP2*, *CD69*, the effector molecule *GZMK* and also the chemokine receptor *CXCR4* (Fig. 3b,c), previously observed in ineffective T-cell responses in cancer^44,45^.

**Fig 3.**
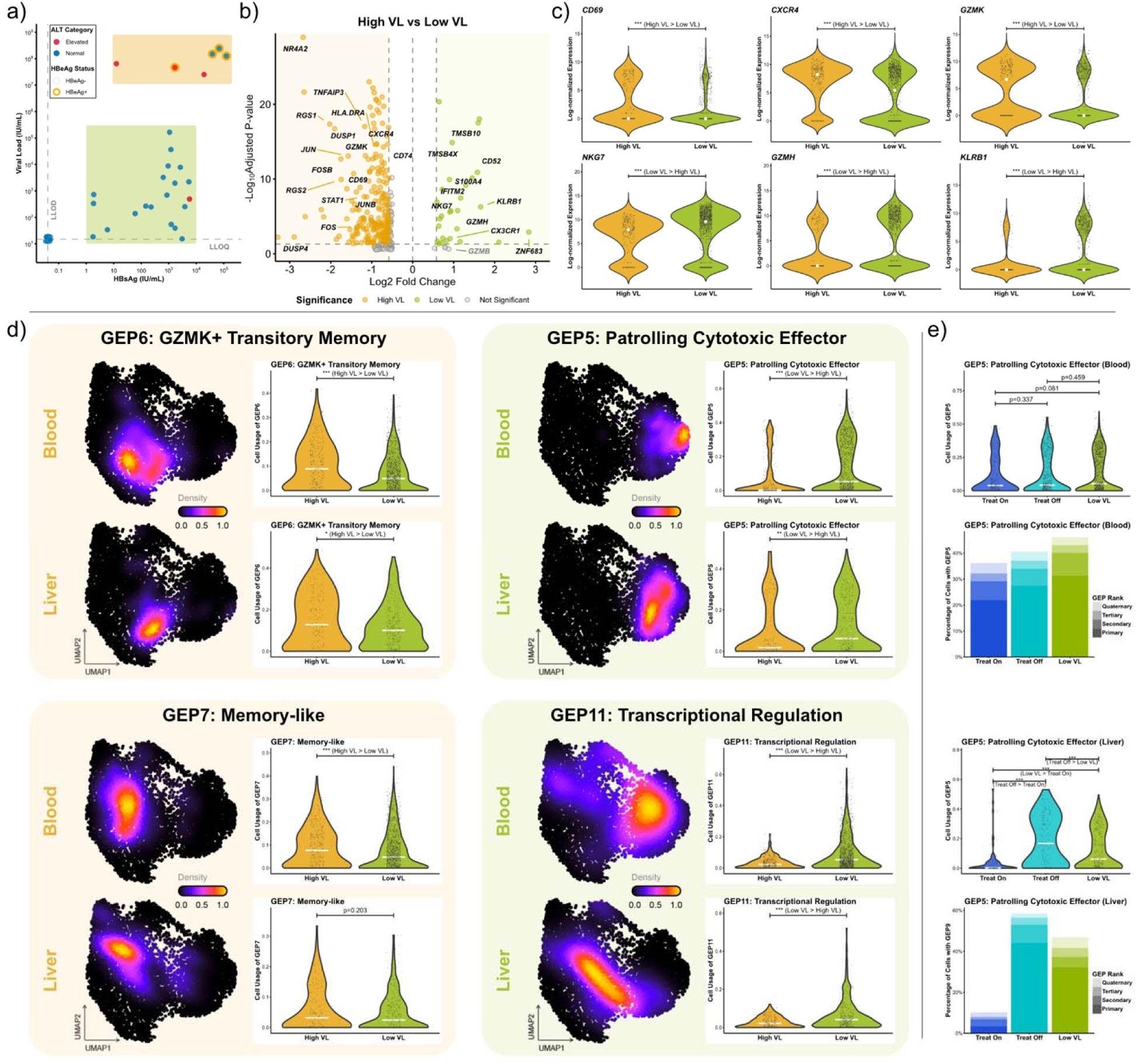
Canonical cytotoxic T-cell signature in control of hepatitis B viral load. (**a**) Distribution of participants along their HBV viral load versus HBsAg. CHB participants with high viral load (orange) are compared with low viral load (green). ALT category (elevated in red, normal in blue) and HBeAg status (orange circle is positive) are indicated (n = 38 participants). (**b**) Volcano plot depicts DGE between blood-derived cells from the respective cohorts (1775 cells, n = 27). Genes of interest with > 1.5-fold change and adjusted *p*-value < 0.05 are labeled. (**c**) Violin plots show expression of selected genes in blood-derived HBV-specific CD8 T cells from the respective cohorts. (**d**) Significant GEP enriched in high viral load (left panel, n = 6) or low viral load (right panel, n = 21) are indicated. UMAP shows density of cells with high GEP usage for blood and liver. Violin plots show selected GEP usage between cells from High VL and Low VL. (**e**) Violin plots show *Patrolling Cytotoxic Effector* GEP usage of blood-derived and liver-derived cells between CHB on long-term treatment (Treat On, blue), after treatment cessation (Treat Off, cyan), and (treatment-naive) Low VL (green). Hierarchical stacked bar chart shows abundance of cells with the *Patrolling Cytotoxic Effector* GEP as their primary highest usage, secondary, tertiary, or quaternary GEP. Statistical significance was determined using Wilcoxon rank-sum test (unpaired, nonparametric, two-sided) and for DGE supplemented with Benjamini-Hochberg correction for multiple testing; \**p* ≤ 0.05, ** *p* ≤ 0.01, *** *p* ≤ 0.001, non-significant *p*-value is given if *p* > 0.05.

cNMF analysis allowed us to define GEPs governing HBV-specific CD8 T cells beyond individual differentially expressed genes. In both blood and liver, low HBV VL was associated with increased activity of a coordinated cytotoxic effector GEP, hereafter referred to as *Patrolling Cytotoxic Effector* (GEP5-*GZMB*, *GNLY*, *PRF1*) (Fig. 3d, Extended Data Fig. 4) that extended beyond the upregulated genes identified by DGE above. In addition to the cytotoxic molecules and genes regulating T-cell cytotoxicity, including antibody-dependent cellular cytotoxicity (ADCC), this GEP is characterized by molecules of advanced differentiation (*KLRG1*, *TBX21*, *PRDM1*, *ID2*) and expression of markers typical for patrolling T cells, tissue residency, and T-cell egress (*CX3CR1*, *S1PR5*, *ZEB2*, and *ZNF683* encoding HOBIT). Cells dominated by this effector GEP were localized within a large subregion of the cytotoxic effector cluster (Fig. 3d, compare to Fig. 1c). Conversely, high viral load CD8 T cells were dominated by two distinct functional GEPs, both primarily localized in the GZMK cluster, one representing *Central Memory* (GEP8-*CCR7*, *SELL*, *LEF1*) and the other of *GZMK+ Transitory Memory* (GEP6-*GZMK*, *CCL4*, *EOMES*) T cells and other minor GEPs (Fig. 3d, Supplementary Fig. 3a).

Notably, while *Patrolling Cytotoxic Effectors* (GEP5) were equally rare in CHB with high VL with or without ALT elevation, enrichment of *GZMK+ Transitory Memory* (GEP6) and *Activation* (GEP3-*CD38*, *HLA-DRA*, *HAVCR2*) was found in the setting of inflammation (Supplementary Fig. 4). These data identify distinct functional T-cell effector profiles associated with different levels of HBV viral load, most notably a profile of *Patrolling Cytotoxic Effector* CD8 T cells enriched in the blood and liver of individuals with well-controlled HBV replication.

Having access to longitudinal samples from participants on long-term antiviral therapy who subsequently stopped treatment allowed us to assess the activity of GEPs longitudinally after the end of extrinsic drug-mediated control of HBV replication (Supplementary Table 5). At baseline, GEPs of HBV-specific CD8 T cells in the blood of treated participants were similar to those from CHB with intrinsically controlled HBV viremia, with surprisingly minor changes after treatment cessation (Fig. 3e, Supplementary Fig. 5a). The situation in the liver, however, was different (Fig 3e, Supplementary Fig. 5b), most notably significantly reduced activity of the *Patrolling Cytotoxic Effector* T-cell GEP (GEP5) that is associated with intrinsic control of HBV replication, suggesting that treatment-mediated control of HBV-rendered cytotoxic activity at the site of infection less necessary (Fig. 3e). Four weeks after treatment termination, the intrahepatic T-cell profile had shifted to a dominance of the *Patrolling Cytotoxic Effector* T-cell GEP (GEP5), while viral replication remained controlled in the absence of NA therapy (Fig. 3e). This initiation of the *Patrolling Cytotoxic Effector* T-cell GEP (GEP5) as drug levels wane further supports a critical role for HBV-specific CD8 T cells with the *Patrolling Cytotoxic Effector* program in controlling HBV replication.

### HBV-specific CD8 T-cell populations with an *NK-like* GEP in control of HBsAg

Having identified *Patrolling Cytotoxic Effectors* as the key GEP in individuals who control HBV replication, we next explored whether functional cure of CHB, i.e. HBsAg loss, was associated with differential gene signature and an additional GEP activity in HBV-specific CD8 T cells by comparing CHB participants with controlled viral loads of less than 1,000 IU/mL but detectable HBsAg to participants who had achieved HBsAg loss (Fig. 4a).

**Fig 4.**
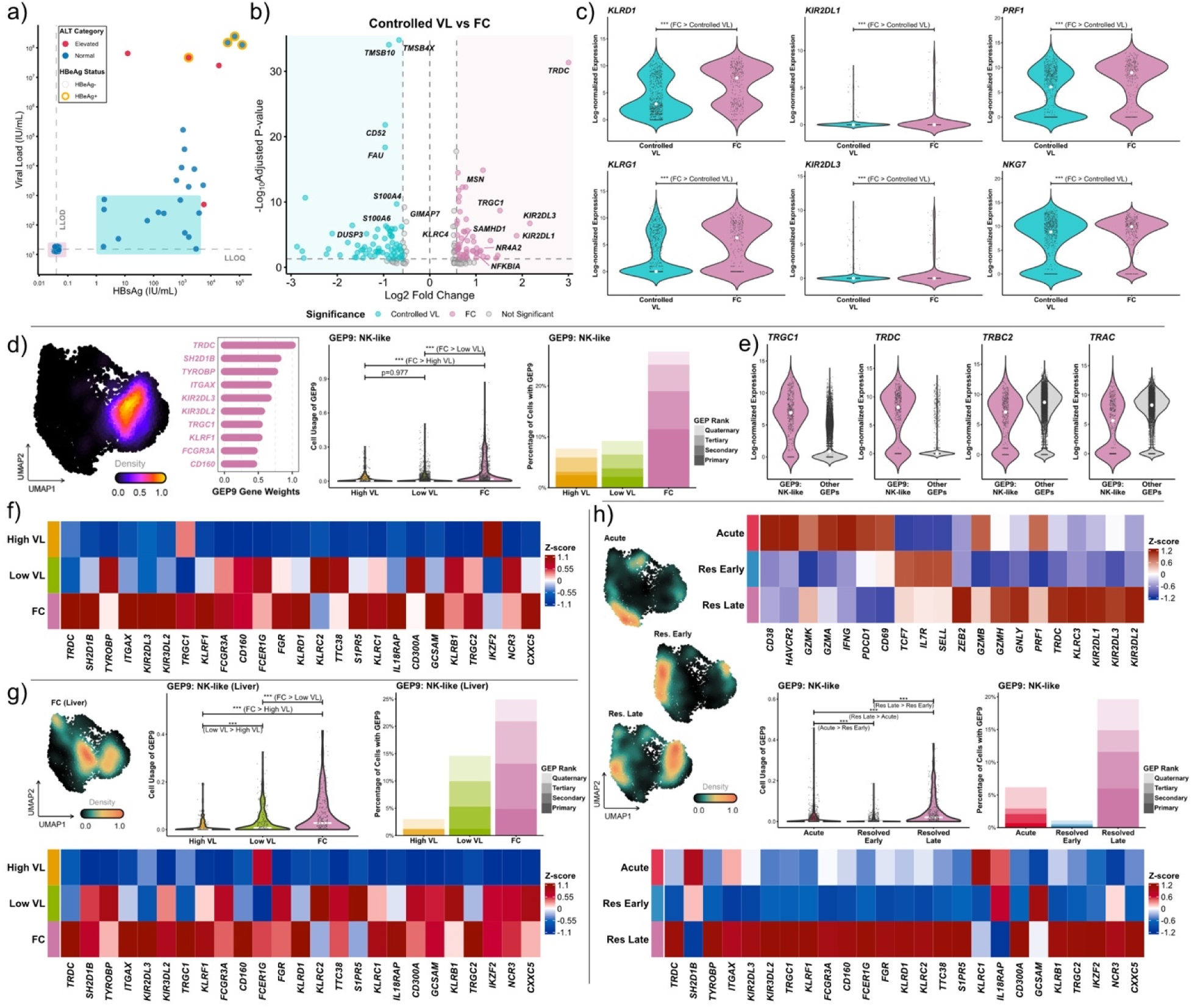
NK-like cytotoxic T cells in control of HBsAg. (**a**) Distribution of participants along their HBV viral load versus HBsAg. CHB participants with controlled viral load (≤ 1000 IU/mL, normal ALT; blue-green) are compared with FC (purple). ALT category (elevated in red, normal in blue) and HBeAg status (orange circle is positive) are indicated (n = 38 participants). (**b**) Volcano plot depicts DGE between blood-derived cells from the respective cohorts (1914 cells, n = 26). Genes of interest with > 1.5-fold change and adjusted *p*-value < 0.05 are labeled. (**c**) Violin plots show expression of selected genes in blood-derived HBV-specific CD8 T cells from Controlled VL (n = 14) and FC (n = 12). (**d**) UMAP shows density of cells with high *NK-like* GEP usage. Horizontal bar graph shows top genes and gene weights defining the *NK-like* GEP. Violin plot shows *NK-like* GEP usage of blood-derived cells between CHB with high VL (yellow, n = 6), low VL (green, n = 21) and FC (purple, n = 12). Hierarchical stacked bar chart shows abundance of blood-derived cells with the *NK-like* GEP as their primary highest usage, secondary, tertiary, or quaternary GEP. (**e**) Violin plots show expression of TCR alpha, beta, gamma, and delta genes in blood-derived HBV-specific CD8 T cells with *NK-like* GEP as primary and secondary usage versus all other blood-derived cells. (**f**) Heatmap with gene expression of top25 genes of *NK-like* GEP of blood-derived HBV-specific CD8 T cells in the indicated cohorts. Gene expression is indicated as z-score. (**g**) UMAP shows density of liver-derived HBV-specific CD8 T cells from CHB with controlled VL and FC. Violin plot shows *NK-like* GEP usage of liver-derived cells between CHB with controlled VL (blue-green, n = 7) and FC (purple, n = 10). Hierarchical stacked bar chart shows abundance of liver-derived cells with the *NK-like* GEP as their primary highest usage, secondary, tertiary, or quaternary GEP. Heatmap with gene expression of top25 genes of *NK-like* GEP of liver-derived HBV-specific CD8 T cells in the indicated cohorts. Gene expression is indicated as z-score. (**h**) Panel shows HBV-specific CD8 T cells from participants with acute hepatitis B in three stages: during Acute infection (n = 8), Resolved in an early state (<12 months, n = 4), Resolved in a late state (≥ 12 months, n = 4). UMAP shows density of blood-derived HBV-specific CD8 T cells from the indicated cohorts. Violin plot shows *NK-like* GEP usage of blood-derived cells between Acute (red), Resolved Early (blue), and Resolved Late (purple) infection. Hierarchical stacked bar chart shows abundance of blood-derived cells with the *NK-like* GEP as their primary highest usage, secondary, tertiary, or quaternary GEP. Heatmap with gene expression of top25 genes of *NK-like* GEP of blood-derived HBV-specific CD8 T cells in the indicated cohorts. Gene expression is indicated as z-score. Statistical significance was determined using Wilcoxon rank-sum test (unpaired, nonparametric, two-sided) and for DGE supplemented with Benjamini-Hochberg correction for multiple testing; \**p* ≤ 0.05, ** *p* ≤ 0.01, *** *p* ≤ 0.001, non-significant *p*-value is given if *p* > 0.05.

DGE analysis between HBV-specific CD8 T cells from the blood revealed increased expression genes typically associated with NK or other innate immune cells, including various KIRs and KLRs, as well as T-cell receptor gamma and delta chains (TRDC, TRGC1) in participants who achieved HBsAg loss (Fig. 4b,c, Supplementary Table 4). These genes are typically associated with human NK cells, including TRGC1 and TRDC which have surprisingly high expression in human NK populations^46,47^. Based on the coordinated expression of these genes within one GEP, we refer to as *NK-like* (GEP9-*TRDC*, *TYROBP*, *KIR2DL3*), with much higher activity in HBsAg loss, that also included the transcription factors *IKZF2* (encoding HELIOS), *ZBTB16*, and *TBX21* (encoding T-BET) (Fig. 4d, Supplementary Table 3). Cells dominated by this GEP were located in the GZMB cluster, in a distinct area from the *Patrolling Cytotoxic Effector* CD8 T cells (GEP5) associated with control of HBV replication and also expressed cytotoxic markers (Fig. 4d, compare with Fig. 1c,d). This *NK-like* GEP was somewhat unexpected and unusual given that we analyzed index-sorted CD3+ CD8+ multimer+ cells that were negative for the NK-cell marker CD56 on the protein-level. Importantly, cells with this GEP also expressed the alpha and beta TCR chains in addition to *TRDC* and *TRGC* (Fig. 4e), a hybrid gene expression profile that has been reported previously before^48^. Nevertheless, there are few reports in the literature for HLA-peptide multimer-positive virus-specific CD8 T cells with an *NK-like* gene expression profile. Expression of the majority of the top genes of GEP9 was higher in FC than in cells from CHB with High VL or Low VL (Fig. 4f). Importantly, enrichment of this *NK-like* program (GEP9) was also observed in HBV-specific CD8 T cells from the liver of participants who achieved HBsAg loss (Fig. 4g).

In addition, adult-acquired acute HBV infection – with its typical spontaneous and complete control of HBV infection – also resulted in a similar CD8 T-cell phenotype to that observed in the blood and liver of CHB with HBsAg loss. In this participant group, we only had access to blood samples, but we were able to assess HBV-specific CD8 T cells longitudinally from active HBV infection through the early phase of HBV control to the long-term HBsAg-negative state more than a year after resolution of acute infection (Fig. 4h, Supplementary Fig. 6, Supplementary Table 6). In these participants, we initially observed differentiation of the HBV-specific CD8 T-cell state with markers of activated effector T cells (*CD38*, *PDCD1*, *IFNG*) and two GEPs reflecting *Activation* (GEP3) and *Active Proliferation* (GEP2) in the acute, HBsAg-positive phase towards a profile resembling classical *Memory-like* (GEP7) and *Central Memory* (GEP8) (*IL7R*, *TCF7*) early after HBV control. However, this memory profile evolved further, as extended control of HBV was associated with the upregulation of cytotoxic (*GZMB*, *GZMH*, *GNLY*) and innate genes (*KLRC3*, *KIR2DL3*, *KIR3DL2*) and the same *NK-like* GEP (GEP9-*TRDC*, *TYROBP*, *KIR2DL3*) as in FC (Fig 4h).

Together, these data from three independent analyses suggest a role of *NK-like*, virus-specific CD8 T cells in the long-term control of HBV, irrespective of whether full HBV control develops after acute or chronic infection. The data from acute infection, where this characteristic GEP emerged only many months after HBV control, further suggest that this novel CD8 T-cell profile might be less relevant for active viral clearance and more involved in long-term control of the remaining viral reservoir.

### Transcriptional control of classical and NK-like cytotoxic GEPs

In order to better define the transcriptional regulation and shared features of the *Patrolling Cytotoxic Effector* and the *NK-like* cytotoxic GEPs that were associated with greater control of HBV replication and of HBsAg, respectively, we defined their characteristic transcription factors (TF) and their correlation with expression of other leading genes in each GEP (Fig. 5, Supplementary Fig. 7). To that end, we employed inferred Gene Regulatory Networks between TF identified within the GEP and the top genes in the respective GEP over the expression within HBV-specific CD8 T cells of CHB participants with high VL, low VL, or functional cure. We observed shared expression of the transcription factor *TBX21* (encoding T-BET) in *Patrolling Cytotoxic Effector* (GEP5) and the *NK-like* GEP (GEP9), which correlated with the expression of cytotoxic molecules such as *GZMB*, *NKG7*, and *PRF1*, consistent with both GEPs being enriched in the GZMB cluster. Its network activity was high in both CHB with low VL and FC. The more canonical *Patrolling Cytotoxic Effector* program (GEP5) and its functional and T-cell mobility genes were also strongly associated with expression of the transcription factors *ZEB2* and *PRDM1* (encoding BLIMP-1). By contrast, the innate-like features of the *NK-like* GEP (GEP9) correlated with the transcription factors *IKZF2*, *CXXC5*, *IRF8*, and *ZBTB16* (encoding PLZF), all of which had up-regulated network activity in FC.

**Fig 5.**
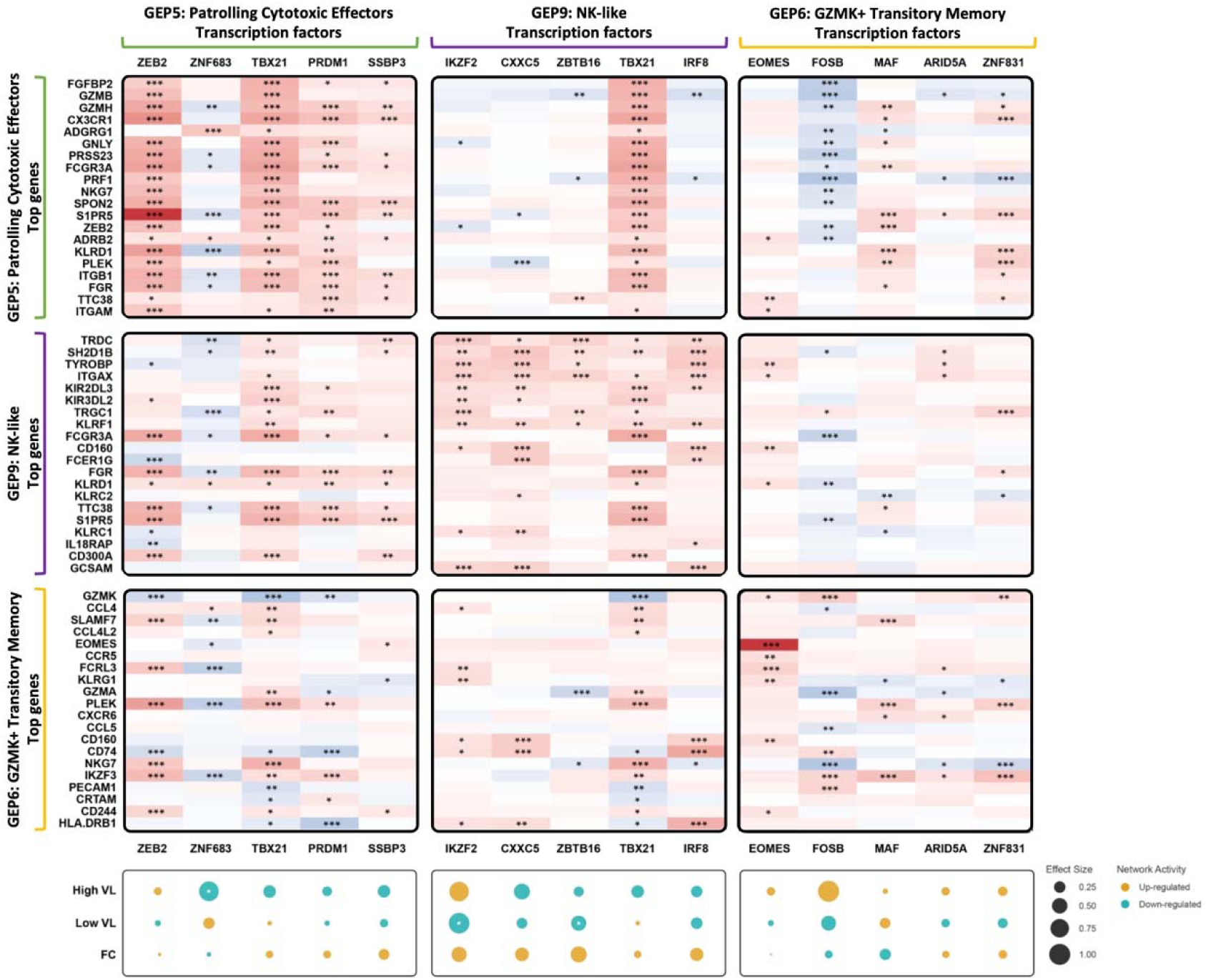
Transcription factor enrichment of different cytotoxic T-cell programs. Heatmap with correlation between top5 transcription factors (x-axis) of three effector GEPs *Patrolling Cytotoxic Effector* (left), *NK-like* (middle), and *GZMK+ Transitory Memory* (right), with top20 genes (y-axis). Dotplot shows network activity according to Gene Regulatory Network for the indicated TF within the GEP for HBV-specific CD8 T cells from CHB participants with High VL, Low VL, and FC. Effect size is indicated by the size of the dot. The effect indicates either an up-regulation (yellow), or a down-regulation (teal). Correlation was determined using Spearman’s rank correlation. Statistical significance was determined using Wilcoxon rank-sum test (unpaired, nonparametric, two-sided) with Benjamini-Hochberg correction for multiple testing; \**p* ≤ 0.05, ** *p* ≤ 0.01, *** *p* ≤ 0.001, non-significant *p*-value is given if *p* > 0.05.

These TF are also active in other innate-immune-cell populations specifically plasmacytoid dendritic cells (pDC), the development of monocytes, and NK cells differentiation^49–54^. In comparison, *GZMK+ transitory memory* (GEP6) had completely distinct transcriptional drivers and gene expression associations underscoring the fundamentally different differentiation states of T-cell populations dominated by this program. This was reflected by the up-regulation of the gene network of all the GEP6-related TF (including *EOMES*) in CHB with high VL. This analysis suggests a shared transcriptional core of cytotoxic capabilities in GEP5 and GEP9, but also a unique transcriptional axis driving innate-like regulation in the *NK-like* GEP.

### *NK-like* HBV-specific CD8 T cells are distinct from KIR-expressing CD8 T cells described in autoimmunity and other viral infections

CD8 T cells expressing KIRs and other innate markers have been described in mice and in humans, mostly in the setting of autoimmunity and some viral infections^55–59^. In these settings KIR+ T cells differed from typical polyclonal antigen-specific CD8 T cells through a narrow TCR repertoire and regulatory cell features. Aiming to establish whether the HBV-specific CD8 T cells we identified in HBV FC are closely related to these regulatory KIR+ CD8 T-cell populations, we utilized a previously published, extensive human gene expression dataset from autoimmunity and COVID-19^58^ for an integrated cNMF analysis also containing our HBV data (Fig. 6a). Within the GEPs defined in the combined data, two GEPs were dominating most of the cells sorted in the autoimmunity/COVID-19, as non-specific KIR+ CD8 T cells, but both GEPs were only lowly detected in HBV-specific CD8 T cells (Fig. 6a). Conversely, another GEP in this combined data set resembling the novel *NK-like* GEP (GEP9) was almost exclusively expressed by HBV-specific CD8 T cells in CHB with HBsAg loss. This analysis demonstrates that *NK-like* HBV-specific HLA-peptide multimer-positive CD8 T cells found in complete HBV control are distinct from regulatory KIR+ CD8 T cells thought to be involved in peripheral tolerance.

**Fig 6.**
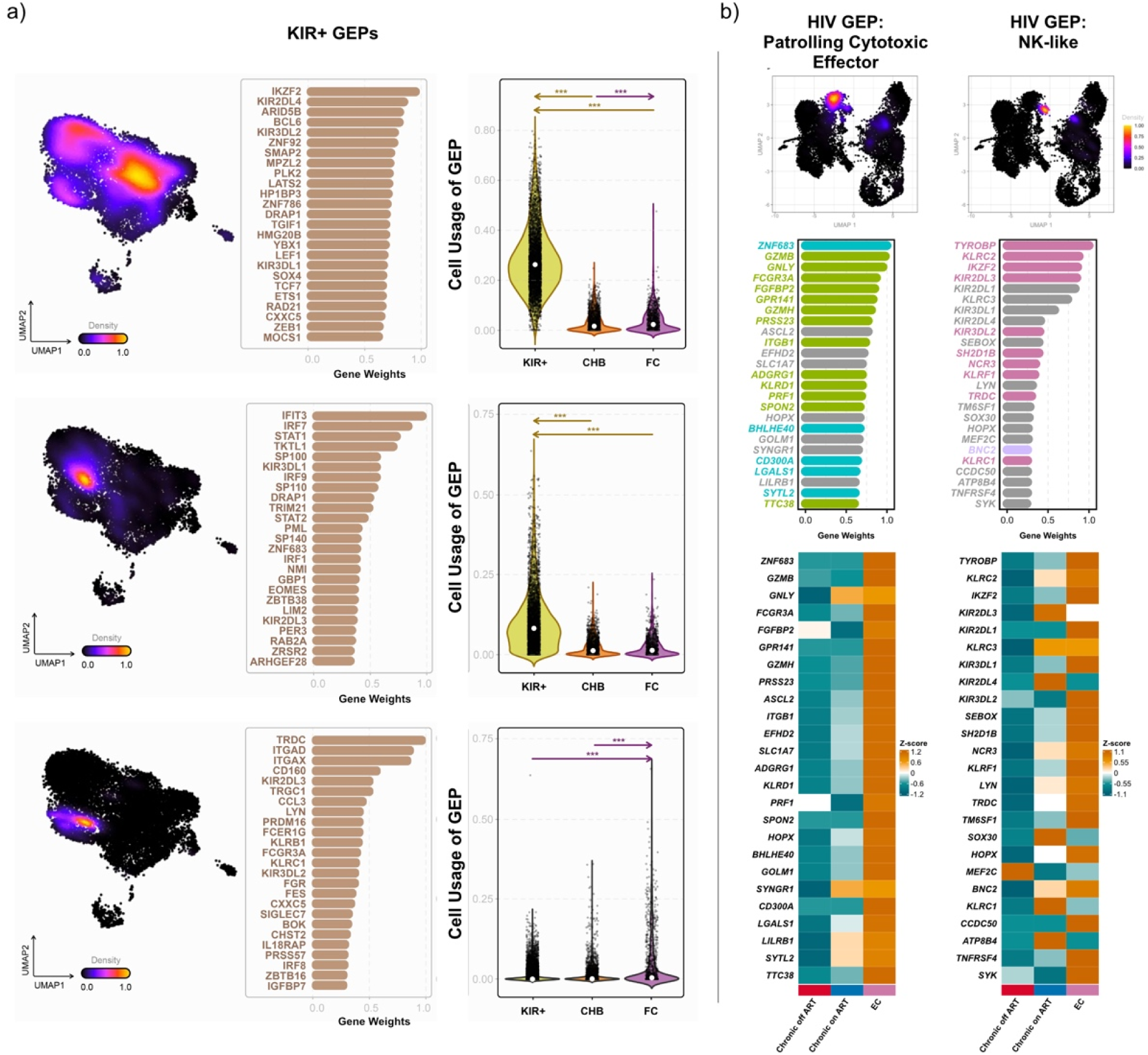
NK-like T-cell signatures in other disease settings. (**a**) Transcriptomes of sorted KIR+ CD8 T cells from Li *et al.* 2023 were integrated with HBV-specific CD8 T-cell transcriptomes^58^. Three GEPs similar to the herein described *NK-like* GEP were identified. UMAP shows density of cells with high GEP usage. Horizontal bar graph shows top genes and gene weights defining the respective GEP. Violin plots show selected GEP usage between cells from the external dataset of KIR+ cells, CHB, and FC. (**b**) Transcriptomes of sorted HIV-specific CD8 T cells from individuals with chronic HIV on-or off-treatment and exceptional and elite controllers of HIV (EC) were analyzed. Data of two selected GEPs are visualized. UMAP shows density of cells with high GEP usage. Horizontal bar graph shows top genes and gene weights defining the GEP. Genes overlapping between the GEP and from the HBV-specific described *Patrolling Cytotoxic Effector* GEP (green in overlap within top25 genes, cyan in top100 genes) and *NK-like* GEP (pink in overlap within top25 genes, purple in top100 genes) respectively, are highlighted. Heatmap with gene expression of top25 genes of *Patrolling Cytotoxic Effector* GEP and *NK-like* GEP of HIV-specific CD8 T cells in the indicated cohorts. Statistical significance was determined using Wilcoxon rank-sum test (unpaired, nonparametric, two-sided); \**p* ≤ 0.05, ** *p* ≤ 0.01, *** *p* ≤ 0.001, non-significant *p*-value is given if *p* > 0.05.

### Classical and *NK-like* cytotoxic programs in HBV-specific CD8 T cells converge with HIV-specific CD8 T cells from exceptional HIV controllers

Elite HIV control is characterized by durable suppression of plasma viremia to undetectable levels due to a functional and specific CD8 T-cell response^60^. In exceptional HIV control^6^, there is an absence of detectable full-length integrated provirus or its presence solely at transcriptionally repressed sites. In these rare instances, HIV is undetectable in the blood, similar to HBV in CHB FC or controlled acute HBV infection where traces of HBV DNA can usually only be detected in liver biopsies^4^. We had the opportunity to analyze two scRNA-seq datasets of HIV-specific CD8 T cells from cohorts including elite and exceptional HIV controllers (please see companion manuscript) (Fig. 6b). One of the two HIV cohorts closely resembled our cohort structure as it analyzed HIV-tetramer-sorted cells directly *ex vivo* from participants with chronic HIV on and off treatment (CP-on and CP-off), and from elite controllers (EC). We identified several GEPs closely resembling GEPs defined in HBV-specific CD8 T cells above (Fig. 6b, Extended Data Fig. 6). Notably, these included the *Patrolling Cytotoxic Effector* GEP as well as the *NK-like* program, containing many key genes shared between HIV and HBV GEPs. As in HBV infection, both distinct cytotoxic GEPs were enriched in HIV-specific CD8 T cells from elite controllers (Fig. 6b). A similar pattern was present in the second HIV cohort, in which HIV-specific CD8 T cells from exceptional controllers were analyzed directly *ex vivo* and compared with CD8 T cells from chronic HIV infection on ART and from exceptional controllers after peptide-specific *in vitro* expansion (Extended Data Fig. 6). Together, the data from HIV controllers strongly support our observations regarding HBV-specific CD8 T cells, suggesting convergent cytotoxic CD8 T-cell programs in complete long-term control of chronic viral infections that more commonly manifest with active viral replication and antigenemia.

## Discussion

HBV-specific CD8 T cells are critical for control of HBV infection^61^, yet their specific functions and molecular profiles associated with viral control remain poorly defined. CHB represents a distinct and complex chronic infection, as it evolves over decades from high-level viremia to control of viral replication as well as certain HBV antigens and, in some individuals, complete suppression of both viremia and antigenemia^9,10^. By generating scRNA-seq profiles of the HBV-specific CD8 T-cell compartment from across all clinical stages of HBV infection, we identified transcriptional programs associated with viral control and functional cure in cells from the blood and confirmed them in T cells from the liver as the exclusive site of infection. These findings from HBV-specific CD8 T-cell populations across diverse HLA-restrictions, broad epitope specificities, and two tissue compartments support a novel framework of HBV-specific T-cell differentiation.

Typically, chronic viral infections with high viral loads, such as HCV or HIV infection, are characterized by CD8 T-cell responses that are terminally differentiated and exhausted (PD-1hi CD127-) together with a smaller proportion of PD-1mid CD127+ memory-like cells^36,40,62–64^. While terminally differentiated T cells may have been generated and subsequently deleted earlier during the extremely high viral load phase^65,66^, they were rare across all stages of CHB, even in individuals whose viral loads exceeded 10^7^ IU/mL, and remained uncommon at mid-to-low viral loads including in CHB on antiviral treatment. By contrast, terminally differentiated HBV-specific CD8 T cells that one would expect in a high antigen load setting were only predominant during adult-acquired acute resolving HBV infection. Tex precursors^42,67^, defined as *TCF7*+, *PDCD1*int, *TOX*+ were also rare in CHB and confined to the highest viral load settings. As these memory-like precursors of exhausted T cells are the key population reinvigorated by immune checkpoint inhibitors^30,44^, these data provide a potential explanation for the limited efficacy of PD-1 blockade for CHB^43,68–70^. These results are also congruent with the clinical course of CHB, where control of viral replication steadily increases in most persons^9,10^, in contrast to HCV or HIV infection where persistently high viral replication is associated with terminally exhausted, dysfunctional T-cell populations^11,12^.

CHB’s complexity is displayed by the fact that viremia and antigen load are not as tightly linked as in other chronic viral infections. This is most obvious in individuals with controlled HBV replication, as HBsAg levels can vary greatly whether viremia is suppressed intrinsically or on antiviral treatment^71^. This complexity likely reflects differential immune mechanisms that control viral replication and HBV antigens, respectively, and may explain the heterogeneous immunological data within CHB cohorts. Comparison of CHB individuals with high versus low levels of viral replication, identified a GEP we termed *Patrolling Cytotoxic Effector* enriched during viral control. This program showed coordinated activity of hallmark genes associated with cytotoxicity, effector cell transcriptional control, and T-cell trafficking. Its biological relevance was further supported in intrahepatic and longitudinal analyses. This GEP is downregulated in T cells from the liver of individuals on long-term antiviral therapy, i.e. when viral replication is controlled extrinsically, only to reemerge after treatment withdrawal despite continued control of viral replication. This longitudinal observation was limited to intrahepatic CD8 T cells, indicating that studying immune cells from the liver might be most important during transitions between clinical phases or treatment-associated changes in viral control where T cells circulating in the blood might not immediately mirror the immunological dynamics in the liver. Importantly, this program correlated with viral control, especially when HBsAg was also suppressed, but not consistently with HBsAg levels, fitting the idea that immune mechanisms governing viral replication and antigenemia overlap only partially. These findings suggest that inducing *Patrolling Cytotoxic effectors* may be necessary, but not sufficient for successful finite therapies that induce functional cure.

In individuals who achieved control of both HBV replication and HBsAg, we identified a second and rather unexpected GEP for an analysis of antigen-specific, HLA-peptide multimer-positive CD8 T cells. This program, which we named *NK-like*, was dominated by genes for innate receptors, NK-signaling adapters, and transcription factors typically associated with NK-cell differentiation and maintenance. The *NK-like* GEP was extremely rare in uncontrolled CHB and not observed in other chronic viral infections such as CMV or EBV (data not shown). By contrast, it was enriched in CD8 T cells from three distinct settings of HBsAg control, i.e. circulating in FC, intrahepatic in FC, and circulating following resolution of adult-acquired acute HBV infection. Notably, in acute HBV infection, this GEP emerged only months after full resolution of infection, suggesting a role in maintaining, rather than initiating full HBV control. *NK-like* HBV-specific CD8 T cells share features with the *Patrolling Cytotoxic Effector* CD8 T cells associated with control of replication, including a neighboring localization in the same cytotoxic cluster and a shared transcription factor (*TBX21*). They also are likely to have differentiated from classical antiviral T-cell effectors since longitudinal analysis of the same specificities during acute-resolving infection initially displayed transcriptional profiles of conventional memory before transitioning to the *NK-like* program.

We propose that this novel *NK-like* CD8 T-cell phenotype with innately regulated cytotoxic features may be particularly suited for maintaining long-term HBV control compared to classical CD8 T-cell memory. As HBV persists in a latent reservoir and hepatocytes might activate HBV replication at any time, a rapid surveillance by innately regulated cytotoxic cells could be more efficient than waiting for conventional memory T cells to reactivate and differentiate into effectors. This model parallels the role of tissue-resident innate-like lymphocyte populations such as invariant NKT cells and MAIT cells which provide rapid responses to recurring microbial stimuli. This suggests that HBV immunity may have co-opted an analogous mechanism within its specific CD8 T-cell compartment. The absence of similar cells in previous studies in humans or model infections is not surprising. Most animal models of infection focus on the dichotomy of acute versus chronic infection and thus do not mirror the situation present in containment of a persistent viral reservoir like in CHB FC.

The relevance of virus-specific CD8 T cells with these two cytotoxic GEPs is further supported by data from two independent scRNAseq datasets in HIV infection. HIV-specific CD8 T cells displayed two GEPs closely resembling the *Patrolling Cytotoxic Effector* and *NK-like* programs we identified in control of HBV replication and antigenemia and they were strongly enriched in persons with exceptional HIV control. The co-occurrence of these two GEPs across two distinct chronic infections suggests a generalizable immunological endpoint of successful long-term viral containment and provides a framework for therapeutic vaccine development or adoptive T-cell engineering.

This study has limitations that warrant discussion. First, due to the extremely low frequency of HBV-specific CD8 T cells in CHB, transcriptional analysis required use of the full samples from this multi-site international study, leaving no material for protein analysis via flow cytometry or functional assays. Especially the latter will require substantially larger amounts of PBMC than what can be obtained from standard research blood draws. Robust functional validation will require new cohorts with collection of large volume blood draws or leukapheresis products from participants with CHB FC. In addition, although inclusion of a broad range of diverse HBV specificities expands upon previous studies relying on few epitopes restricted by single HLA alleles, additional unrecognized CD8 responses may have been missed. In the future, defining additional epitopes and simultaneously analyzing dozens of specificities using large DNA-barcoded HLA class I multimer libraries^72,73^ should be able to overcome most of this limitation. Finally, the decades-long natural history of CHB largely necessitated cross-sectional comparisons, limiting our ability to directly observe differentiation of T-cell clones towards these GEP profiles. Ongoing advances in HBV-specific drug treatments that achieve functional cure in a substantial proportion of participants^74^ together with observations from HBV/HIV coinfection where initiation of standard antiviral treatment leads to higher numbers of functional cure^75,76^ create an opportunity for longitudinal studies that define the CD8 T-cell trajectories associated with successful versus unsuccessful attainment of functional cure of CHB.

In summary, our findings reveal novel gene expression programs in virus-specific CD8 T cells that associate with full control of not only HBV, but also HIV infection, suggesting an important role of these cytotoxic CD8 T-cell phenotypes in chronic infections that are rarely fully contained in the absence of antiviral therapy. Further studies on the functionality and regulation of *NK-like* antiviral CD8 T cells will be essential to inform the design of novel immune interventions aimed at achieving durable control of chronic infection in the absence of continued antiviral treatment.

## Methods

### Study design, HBV cohort and sample collection

As part of a multicenter research study we enrolled participants in all untreated stages of CHB, in CHB with HBsAg loss, in CHB on long-term treatment, and with adult-acquired acute HBV infection. Exclusion criteria was advanced fibrosis and coinfection with other hepatitis viruses, or HIV. The clinical sites were Toronto General Hospital (Toronto, Canada), Erasmus Medical Center (Rotterdam, The Netherlands), Massachusetts General Hospital (Boston, USA), Fiocruz (Rio de Janeiro, Brazil), and Janssen Clinical Pharmacology Unit (Antwerp, Belgium). Local ethic committees approved the study and it was conducted in accordance with both the declaration of Helsinki and Istanbul. All participants provided written informed consent. For every participant, PBMC were isolated from blood draws by density gradient centrifugation. Intrahepatic lymphocytes were isolated from liver fine needle aspirates that were obtained in 71% of study participants with CHB. FNA procedure was performed as described previously^77^. Both PBMC and IHL were cryopreserved at the local recruitment site. Clinical characteristics of the participants are described in Extended Data Fig 1a.

Participants with CHB on treatment participated in a prospective study that included stopping antiviral treatment and multiple FNAs for research purposes. Entry requirements were longterm treatment (> 3 years), HBV viral load at LLOQ /LLOD for 2 years prior to screening, negative test for HBeAg, ALT < 1.5x ULN for 1 year, and absence of liver cirrhosis. After stopping treatment, participants were closely monitored for 24 weeks to re-initiate treatment in case of a virological or clinical relapse. Virological relapse was defined by HBV DNA reaching 2000 IU/mL or reversion to HBeAg positive and clinical relapse by the occurrence of ALT > 5x ULN. After treatment re-initiation participants were followed for another 6 months. While 3/5 participants displayed control of HBV replication up to 24 weeks none achieved HBsAg loss and they all relapsed eventually (Supplementary Table 5).

### MHC class I multimers

Pentamers were acquired from ProImmune. MHC monomers were obtained from the National Institute of Health Tetramer Core Facility (Atlanta, USA). Monomers were labeled with phycoerythrin (PE) or allophycocyanin (APC) by tetramerizing 25 µL per monomer at 2 mg/mL µg/mL by adding 25 µg of Streptavidin-APC or Streptavidin-PE in 5 steps with 10 min incubation at 4 °C each. We used multimers for 28 distinct epitopes restricted by 10 different HLA alleles (Supplementary Table 1). Multimers were used at 0.2 mg/mL.

### Screening and staining of virus-specific CD8 T-cell populations

Detection of multimer-specific CD8 T cells was performed both after *in vitro* stimulation (cell lines) and directly *ex vivo* to screen for HBV-specific CD8 T cells as described before^78^. Cryopreserved PBMC were thawed in R10 medium. For the generation of cell lines, PBMC (5 to 10 × 10^6^ cells) were stimulated with single overlapping HBV peptides (10 μg/mL) for 12 days in R10/50 medium (R10 containing recombinant interleukin-2 [IL-2; 50 U/mL; Sigma-Aldrich]). Every other day, fresh R10/50 medium was added. By contrast, *ex vivo* screening was done directly after PBMC thawing and without stimulation.

For the detection of HBV-specific CD8 T cells, cell samples were washed with FACS buffer (PBS + 2% FBS), pelleted, and incubated with a viability dye (Invitrogen LIVE/DEAD Fixable Blue) for 30 min on ice. Cells were washed and pelleted. Cells were incubated with multimers for 10 min at RT, washed, and pelleted. HBV-multimer-positive cells were enriched using MACS columns (Miltenyi Biotec) with anti-PE and anti-APC MicroBeads, according to the manufacturer’s protocol. Cells were then stained with surface antibodies (Supplementary Table 7) for 30 min on ice and either resuspended in 2% paraformaldehyde (Fisher Scientific) in PBS. Compensation and rainbow beads (BD Bioscience) were used for compensation and for detector calibration. Cell samples were acquired on a Becton Dickinson LSR 2 flow cytometer.

84 individuals with CHB or FC were screened for HBV-specific CD8 T cells based on their HLA types (A*0101, A*0201, A*0301, A*1101, A*2402, B*0702, B*0801, B*3501, B*4001, B*5101). Detectable specificities were found in 63% of all participants (58% of individuals with untreated CHB, 60% with treated CHB, and 82% with FC).

### Flow cytometry data analysis and visualization

Flow cytometry data were analyzed using FlowJo software, version 10.10. Protein expression levels were extracted as percentages for gated populations and as individual fluorescence values for each marker. For visualization following FlowJo-style scaling on a biexponential axis, single-cell fluorescence values were transformed with inverse hyperbolic sine.

### Index cell sorting and single-cell RNA-seq library preparation

Singular *ex vivo* HBV-specific CD8 T cells were index-sorted from PBMC and IHL of participants with chronic infection before (n = 27) and after functional cure (n = 12) at the Ragon Institute’s Flow Cytometry Core (Extended Data Fig. 2b). Additionally, longitudinal samples of CHB participants on long-term treatment with treatment interruption (n = 5), or adult-acquired acute (n = 9) infection before and after HBsAg loss in singular wells of 96-well Armadillo plates on a FACS SORP 5-laser (BD Biosciences) (Extended Data Fig. 2c). Cells were sorted with the 100 µm nozzle at 20 psi into RLT buffer + 1% BME buffer. Sample and collection plate were kept at 4 °C during the run. Sorted plates were briefly spun down before freezing on dry ice.

Despite our HBV HLA multimer library covering 10, 9, 7, and 2 epitopes in HBV core, polymerase, envelope, and x, respectively, a majority of the detectable responses and a higher percentage of isolated cells were targeting HBV core, despite its sequence representing less than 20% of the HBV genome. This dominance of detected core responses was observed throughout all disease stages, including FC.

### Single-cell RNA-seq library preparation

Single-cell RNA sequencing and library preparation was performed at the Broad Genomics Platform using the Smart-Seq2 platform^16,17^. Libraries were loaded on a NextSeq500 (Illumina) to generate dual-indexed 38-8-8-38 paired-end reads.

### Sequence Alignment and Gene Quantification

Raw sequencing reads from Smart-Seq2 libraries were aligned to the human reference genome (GRCh38) using HISAT2 (v2.1.0)^79^. Gene-level expression was quantified as both counts and Transcripts Per Million (TPM) using RSEM (v1.3.0)^80^.

### Single-cell RNA-seq Processing and Integration

Data analysis was performed using the Seurat (v5.5.0) package in R^81^. To minimize participant-specific batch effects and focus on biologically conserved signals, we utilized a stratified feature selection strategy. The dataset was split by participant ID, and 2,000 highly variable features were identified within each individual subset using the *vst* method. Consensus variable features were then defined using *SelectIntegrationFeatures*.

High-quality cell states were identified through an iterative filtering and clustering process. An initial unsupervised clustering pass (Leiden, resolution = 1.0) was used to identify and exclude clusters characterized by low gene complexity or technical artifacts. For final dimensionality reduction, we selected eight specific principal components based on Elbow plot analysis to maximize biological variance. Final cell states were defined using the Leiden algorithm (resolution = 0.295) and visualized via UMAP.

### Consensus Non-negative Matrix Factorization (cNMF)

To identify coordinated transcriptional programs, we applied consensus Non-negative Matrix Factorization via the Terra platform^26^. The algorithm factorizes the expression matrix V into a usage matrix W and a gene-weight matrix H: V ≈ W * H, where V is the x*m matrix of cells by genes, W is the n*K usage scores, and H is the K*m gene spectra. We evaluated a range of K values from 2 to 40. After assessing stability and reconstruction error, K=16 was selected. To identify the regulatory drivers of these programs, a final augmented cNMF iteration was executed. In this run, the input gene set was forced to include the 2,000 consensus variable features plus a list of transcription factors. The list of human transcription factors was obtained from the Aerts Lab cisTarget repository, which compiles reference TFs originally defined by Lambert *et al*^82^. This ensured that TF weights were explicitly represented in the gene spectra (H) for every program. Usage scores were normalized per cell for downstream clinical integration.

### Gene Regulatory Network Inference

To identify the co-regulatory architecture driving the transcriptional programs, we implemented a custom inference pipeline. To account for technical sparsity, cells within each clinical and metadata category (participant, timepoint, and antigen) were randomly partitioned into subsets of at least 10 cells. These subsets were subsequently aggregated into pseudobulk samples using mean expression.

For each identified GEP, Spearman’s rank correlations were calculated between the expression of top-weighted transcription factors and the top 50 associated genes. Co-regulatory networks were defined empirically for each TF, consisting of program genes exhibiting a statistically significant correlation (*p* < 0.05) and a minimum coefficient strength of |r| > 0.1.

Network activity scores (a) were calculated for each pseudobulk sample using a custom weighted enrichment scoring pipeline, where the activity of a TF was estimated as:

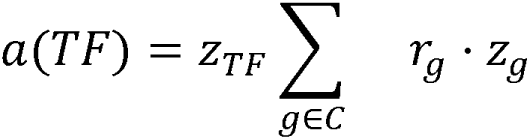

Where C represents the set of co-regulated genes, r_g_ is the Spearman correlation coefficient for gene g, and z_TF_ and z_g_ are the z-score normalized expression values of the TF and its co-regulated genes, respectively. This scoring approach prioritized TFs whose expression was synchronized with the collective intensity of their identified co-regulatory network. Statistical differences in activity scores across clinical cohorts were evaluated using the Wilcoxon rank-sum test, with *p*-values adjusted for multiple testing via the Benjamini-Hochberg method.

### Comparative Analysis and Clinical Integration

Clinical metadata, including HBV VL, HBsAg, and ALT, were integrated with single-cell parameters. Participants were stratified into High VL (≥ 10^6^ IU/mL), Low VL (< 10^6^ IU/mL) and FC groups. Longitudinal samples of resolved acute hepatitis B were categorized according to their time post resolution into “Resolved Early” (≤ 365 days post-resolution) and “Resolved Late” (> 365 days). Differential gene expression was calculated using the Wilcoxon rank-sum test with Benjamini-Hochberg correction. Genes were considered differentially expressed with fold-change ≥ 1.5 (x-axis reflects log2(1.5) = 0.58) and adjusted *p*-value < 0.05.

### Visualization

Gene expression data were visualized via UMAP, DotPlots, Heatmaps, volcano plots and Violin plots. The distribution of programs across clinical categories was visualized using stacked bar charts representing primary, secondary, tertiary, and quaternary usage levels. Sankey diagrams representing cell state transitions and program distributions were produced using the *Flourish* web application. Heatmaps were visualized with *Complex Heatmap*. All other statistical visualizations were generated using *Seurat* and *ggplot2*.

### Statistical analysis

The Wilcoxon-paired test and the Wilcoxon rank sum test (unpaired, nonparametric, two-sided) were used for statistical analysis. For correlations, a Spearman rank correlation coefficient was calculated. Data were summarized using the median and interquartile range (IQR). Significance is displayed as \**p* ≤ 0.05, ** *p* ≤ 0.01, *** *p* ≤ 0.001, non-significant *p*-value is given if *p* > 0.05.

## Data availability

RNA-seq data from this study will be made publicly available through the NCBI Gene Expression Omnibus and/or NCBI database of Genotypes and Phenotypes. The remaining data supporting the findings of this study are available from the corresponding authors upon request.

## End notes

## Acknowledgements

We would like to acknowledge the participants and physicians involved in this study. The NIH tetramer facility for providing the HBV-specific multimers. The University of Oklahoma Medical Center HLA Typing Facility (W. Hildebrand). We thank Nathalie Bonheur of the Ragon Institute’s Flow cytometry core for her advice and assistance. The members of the Ambulatório/ Laboratório de Hepatites Virais, Instituto Oswaldo Cruz/ FIOCRUZ, Rio de Janeiro, Brazil. We thank Paul Klenerman, Jörg Timm, and Jasmin Mischke for critically reading the manuscript. This work was supported by fellowships from the German Research Foundation DFG (to H.K.D. and L.M.B.). Janssen Pharmaceuticals sponsored the study.

## Author Contributions

L.M.B., H.K.D., R.C.H., A.S.G., G.G., A.J.G., and G.M.L. designed the experiments. E.I.L., M.F., L.M.B., H.K.D., S.D., M.A.V., D.L., K.V.D.B., E.D.T., S.S., A.S.G., N.C.-N., A.T., A.A., C.R.C., M.D., J.A.C., M.V.M.-C., and M.W. performed and analyzed experiments. E.I.L., M.F., S.D., D.L., K.V.D.B., E.D.T., and S.S., performed computational analysis. N.C.-N., L.S., B.J.B., D.L., J.D.S.V., St.Sh., N.A., R.J.D.K., L.L.L.-X., J.Ae., J.B., N.H., G.G., R.T.C., J.F., H.L.A.J., A.K.S., A.B., A.J.G., and G.M.L. designed the clinical cohorts and sample acquisition. B.J.B., D.La, J.A., A.Y.K., L.L.L.-X., J.Ae., R.T.C., J.F. contributed to the clinical cohort recruitment and metadata management. M.F., E.I.L., and G.M.L. drafted the manuscript. All authors contributed to and approved the final version of the manuscript. E.I.L. and M.F. contributed equally. G.M.L. was responsible for the overall design and oversight of the experiments.

## Additional Information

Supplementary Information is available for this paper.

Correspondence and requests for materials should be addressed to Georg M. Lauer.

## Extended Data

**Extended Data Fig. 1:**
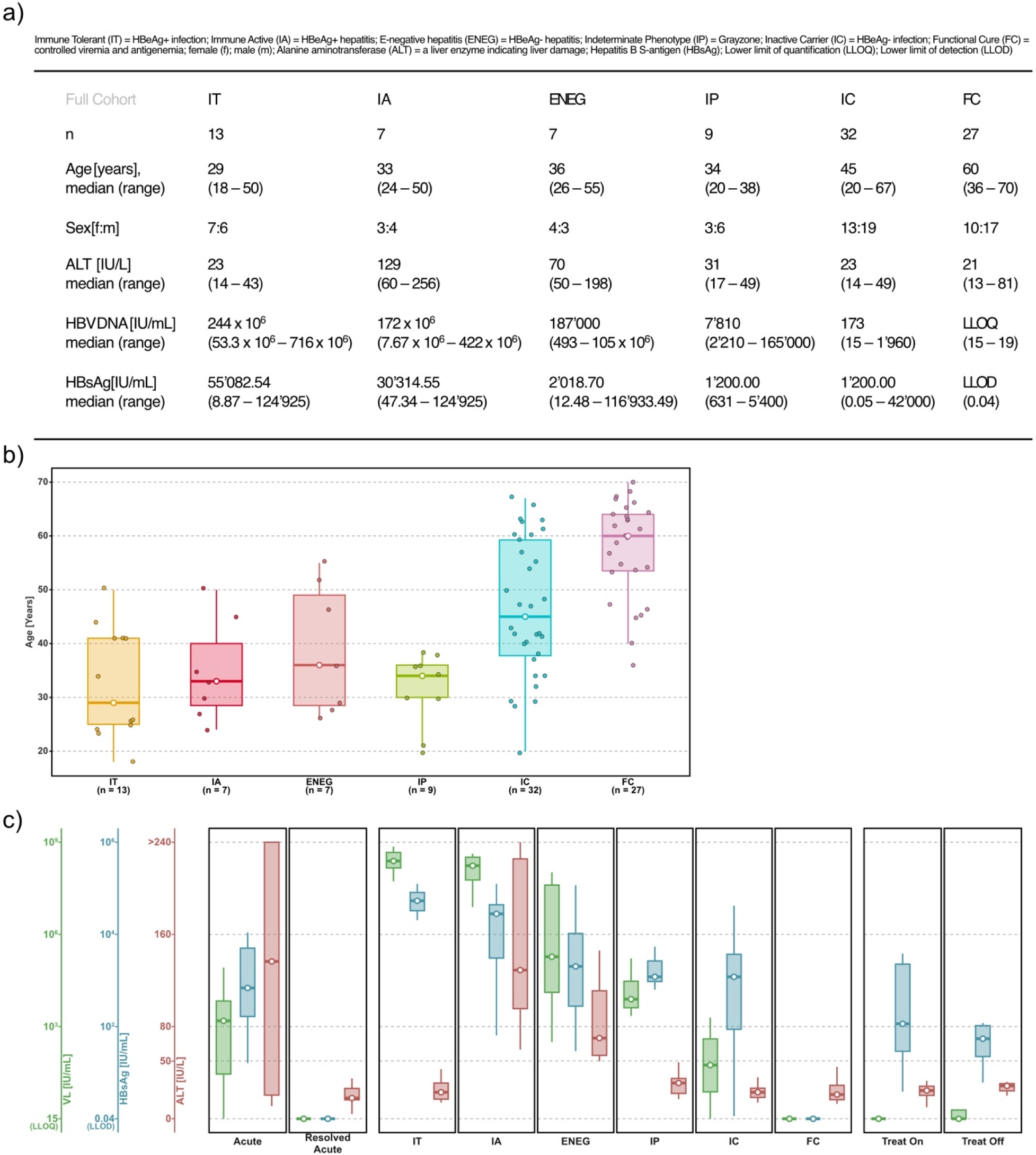
The cohort covers the full breadth of HBV infections. (**a**) Participant overview of consortium-recruited samples. Table indicates number of individuals and their clinical data with median, range, or ratio per cohort as defined by EASL/AASLD guidelines. Immune tolerant (IT) is HBeAg+ infection; Immune active (IA) is HBeAg+ hepatitis; E-negative (ENEG) is HBeAg- hepatitis; Indeterminate phenotype (IP) is Greyzone; Inactive carrier (IC) is HBeAg- infection; Functional cure (FC) is full control of viral replication and antigenemia. (**b**) Age for study participants in the untreated CHB cohorts are given. Dots indicate each participant’s data. Box’s higher and lower limit are indicating interquartile range and white circle is showing the median. Whiskers indicate the range. (**c**) Viral load (VL), HBV S antigen (HBsAg), and liver enzyme Alanine aminotransferase (ALT) are indicated for the full breadth of HBV infections: Acute and Resolved acute infection, the stages of chronic hepatitis B with functional cure, and participants on nucleoside/-tide analogue treatment or with treatment interruption. Box’s higher and lower limit are indicating interquartile range and white circle is showing the median. Whiskers indicate the range.

**Extended Data Fig. 2:**
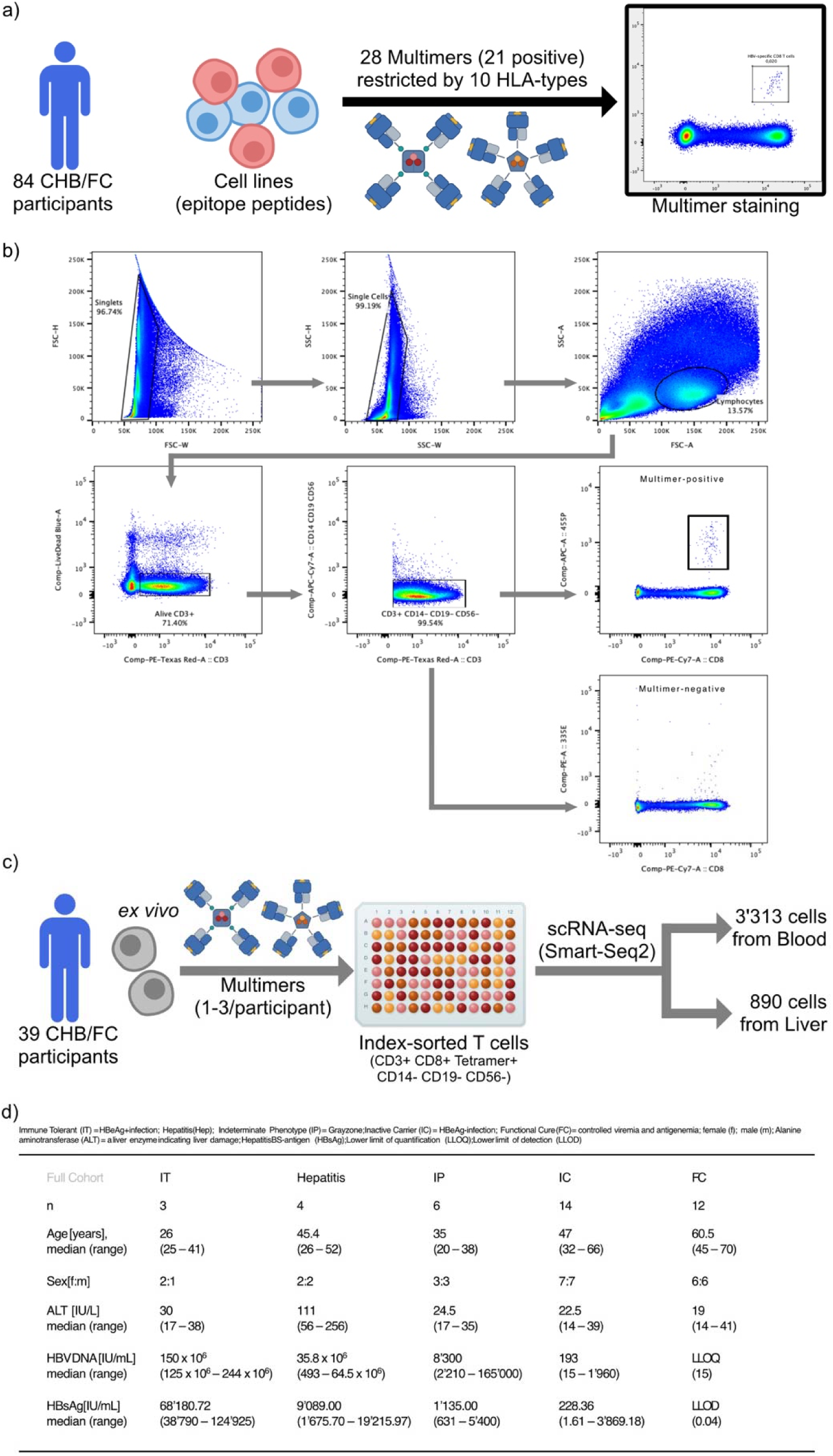
Experimental setup. (**a**) 84 participants with CHB or FC were screened for HBV-specific CD8 T cells. Cells were expanded with 10 days of peptide stimulation and stained with multimers for their respective HLA background. Multimer staining was evaluated with flow cytometry. (**b**) Sorting strategy is depicted by representative flow plots gating singlets, lymphocytes, alive, CD3+, CD14- CD19- CD56-, CD8+ and multimer positive cells. (**c**) Cells (PBMC and/or IHL) of 39 CHB or FC participants with positive multimer staining and sufficient, viable cells were sorted. Cells were not stimulated but stained *ex vivo* with antibodies and 1-3 multimers per participant. Cells were index-sorted based on their individual fluorescence signal. Single cell RNA sequencing of cells in singular wells was performed with Smart-Seq2. (**d**) Overview of participants whose PBMC and/or IHL were sorted. Table indicates number of individuals and their clinical data with median, range, or ratio per cohort.

**Extended Data Fig. 3:**
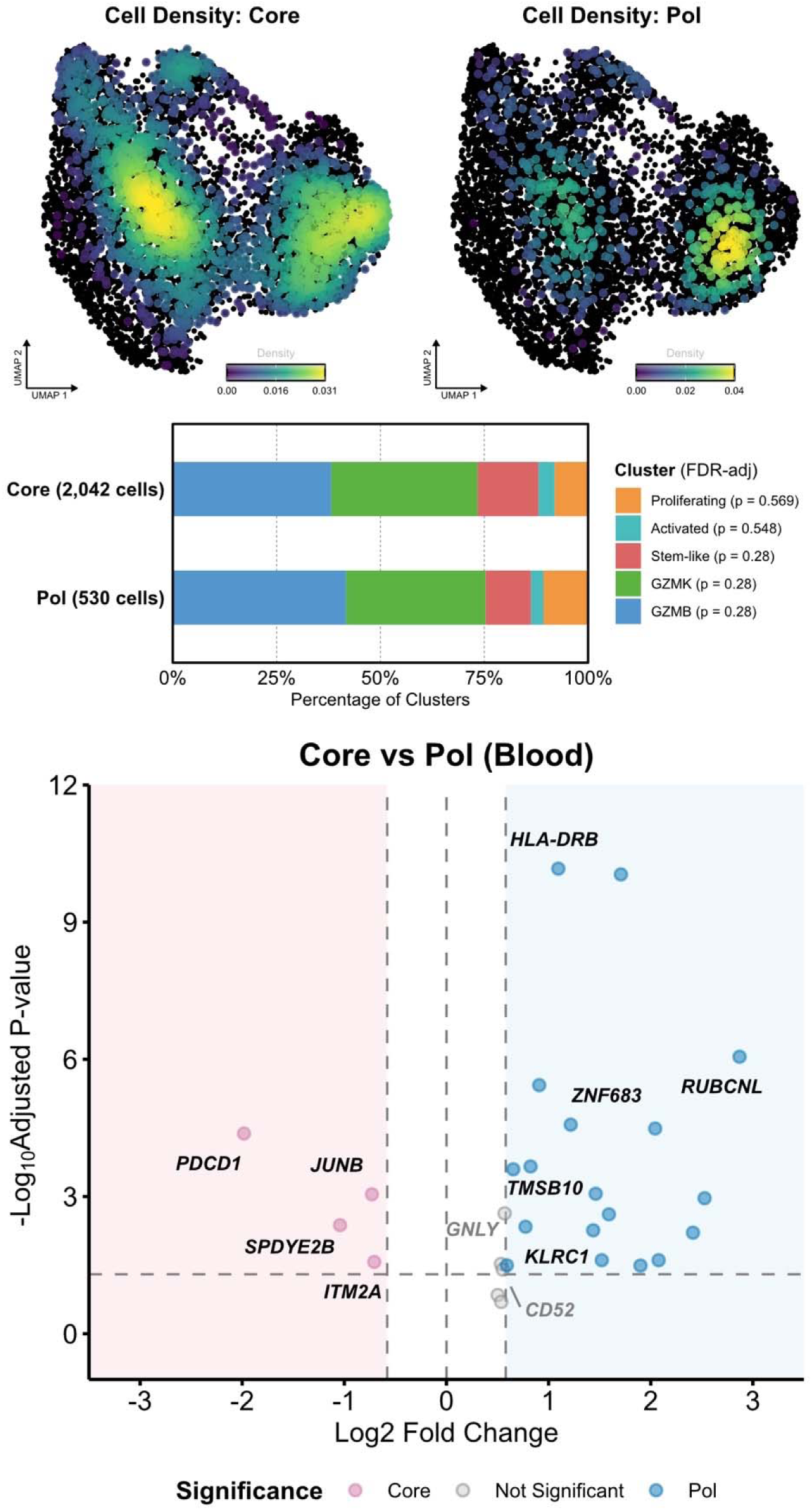
Comparison of Core versus Pol antigen-specific T cells. Distribution of blood-derived HBV Core and Pol-specific CD8 T cells from participants with ongoing chronic hepatitis B (CHB and treated CHB) is shown in the UMAP space. Densely populated areas are indicated in green with the highest density in yellow. Percentage of Leiden clusters (Activated, Proliferating, GZMB, GZMK, Stem-like) in CD8 T cells specific for HBV Core and Pol are shown. Total cell number per antigen is indicated. Significance is shown as a *p*-value in the legend. Volcano plot depicts DGE between blood-derived cells from the respective cohorts (2572 cells, n = 30 participants). Genes of interest with > 1.5-fold change and adjusted *p*-value < 0.05 are labeled (Core in pink, Pol in blue; non-significant in gray). Statistical significance was determined using Wilcoxon rank-sum test (unpaired, nonparametric, two-sided) and for DGE supplemented with Benjamini-Hochberg correction for multiple testing; \**p* ≤ 0.05, ** *p* ≤ 0.01, *** *p* ≤ 0.001, non-significant *p*-value is given if *p* > 0.05.

**Extended Data Fig. 4:**
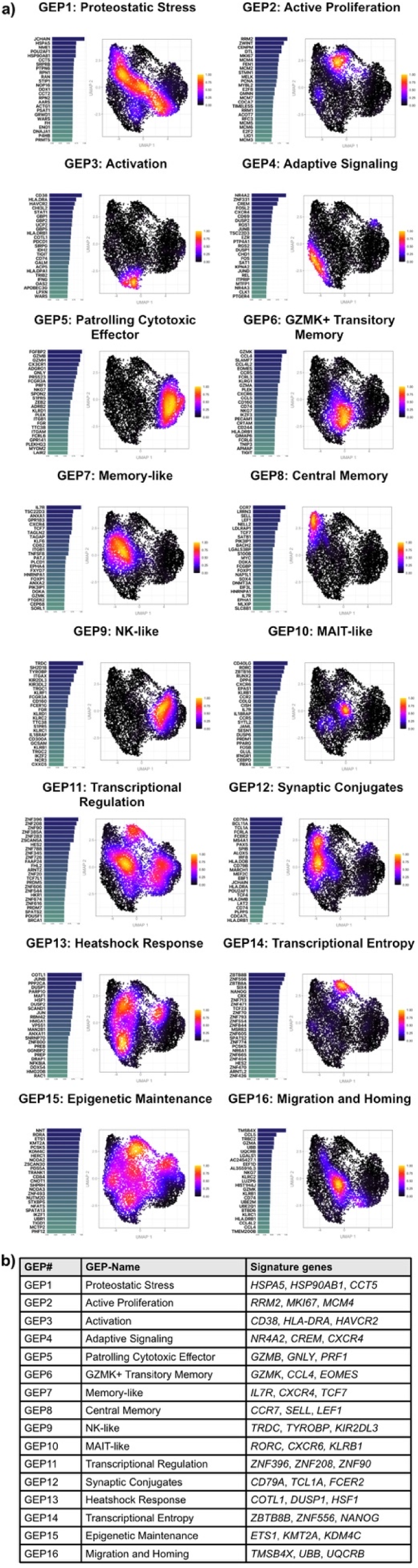
Gene expression program (GEP) overview. (**a**) Horizontal bar graph shows top genes and gene weights defining the 16 GEPs. UMAP shows density of cells with primary GEP usage. (**b**) GEP-number, the respective name, and 3 signature genes are given for the 16 GEPs as determined with cNMF.

**Extended Data Fig. 5:**
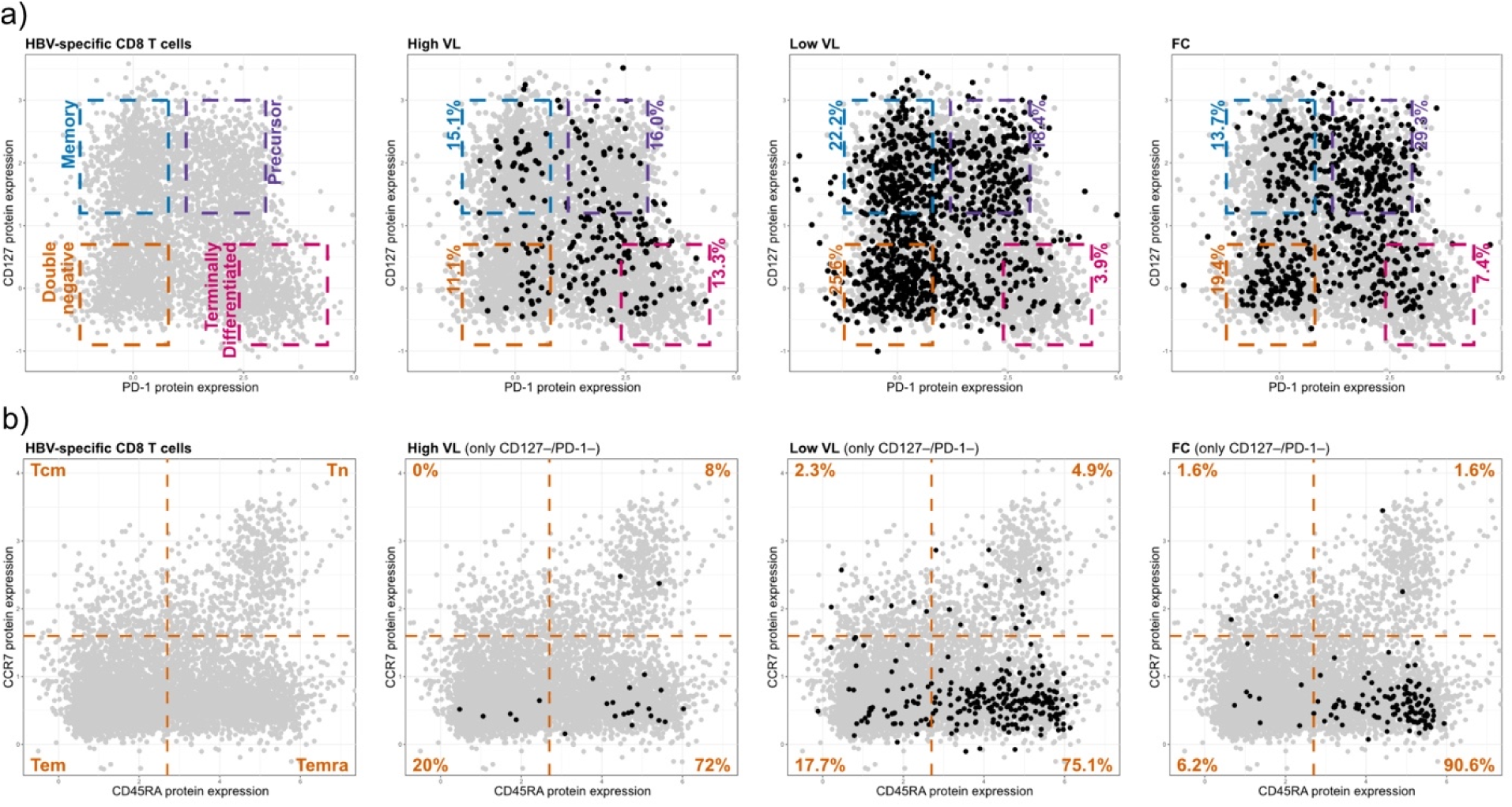
Depletion and exhaustion of HBV antigens. (**a**) CD127 and PD-1 protein data of HBV-specific CD8 T cells (gray) is indicated on the left. Gates of Memory (CD127+ PD-1-; blue), Precursors (CD127+ PD-1 mid; purple), Terminally differentiated (CD127- PD-1hi; pink), and Double negative (CD127- PD-1-; orange) are shown for the respective cohorts. Protein data of HBV-specific CD8 T cells in the indicated cohort (black) versus all HBV-specific CD8 T cells (gray). (**b**) CD45RA and CCR7 protein data of HBV-specific CD8 T cells (gray) is indicated on the left. Gates of naïve (Tn; CD45RA+ CCR7+), central memory (Tcm; CD45RA- CCR7+), effector memory (Tem; CD45RA- CCR7-), and effector memory RA+ (Temra; CD45RA+ CCR7-) T cells are shown for the respective cohort. Protein data of CD127- PD-1- (double negative) HBV-specific CD8 T cells in the indicated cohort (black) versus all HBV-specific CD8 T cells (gray) are shown. Percentage indicates the ratio of black cells in the respective gate of all black cells.

**Extended Data Fig. 6:**
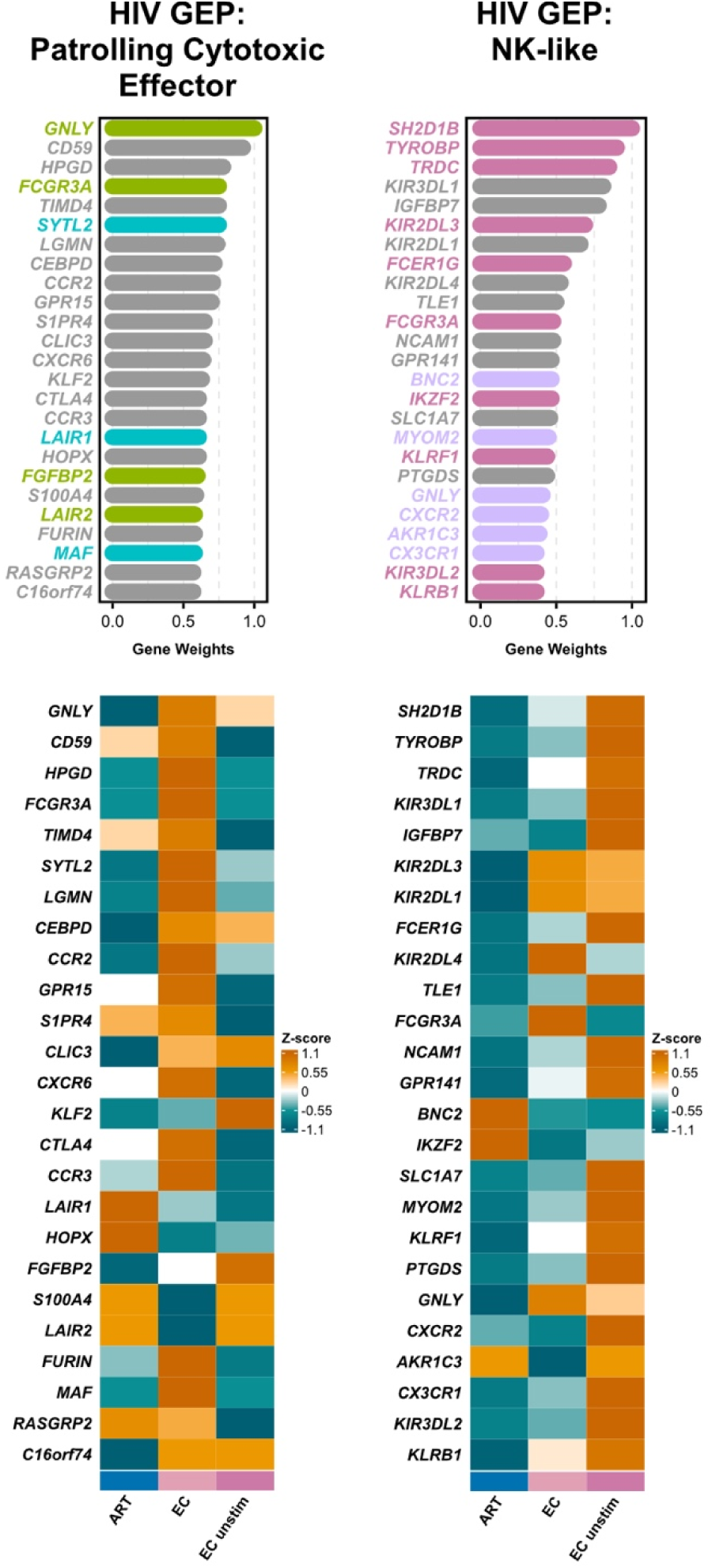
Stimulation of cytotoxic HIV-specific T cells. Transcriptomes of sorted HIV-specific CD8 T cells that were stimulated with HIV-derived peptides. Cells from chronic HIV on-treatment (ART) and exceptional and elite controllers of HIV (EC) were analyzed and compared to unstimulated cells from EC. Data of two selected GEPs are visualized. Horizontal bar graph shows top genes and gene weights defining the GEP. Genes overlapping between the GEP and from the HBV-specific described *Patrolling Cytotoxic Effector* GEP (green in overlap within top25 genes, cyan in top100 genes) and *NK-like* GEP (pink in overlap within top25 genes, purple in top100 genes) respectively, are highlighted. Heatmap with gene expression of top25 genes of *Patrolling Cytotoxic Effector* GEP and *NK-like* GEP of HIV-specific CD8 T cells in the indicated cohorts.

