## Supplementary Figures for "CD8 T cells with classical and NK-like cytotoxic gene expression programs mediate control of HBV replication and functional cure"

Supplementary Material

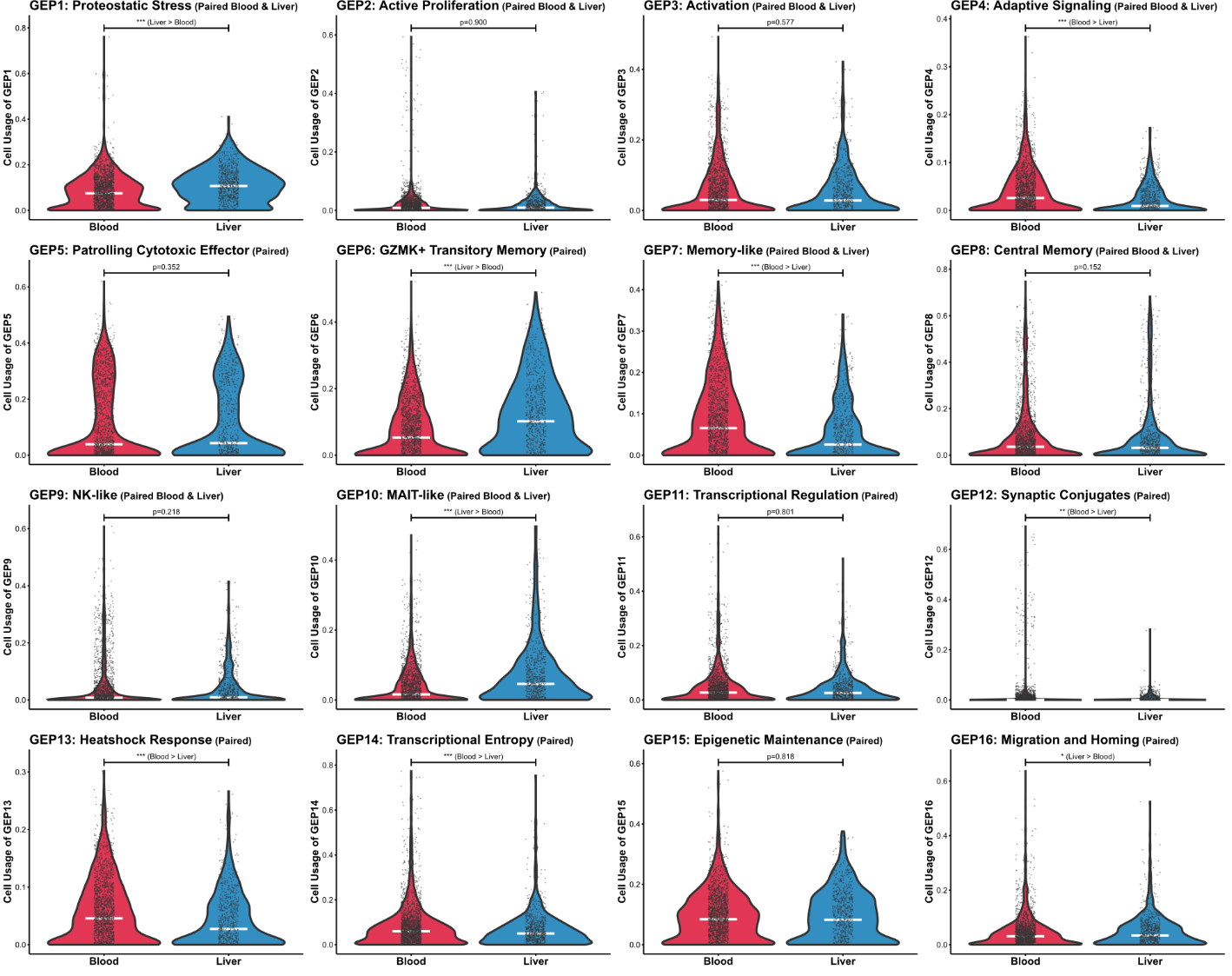

**Supplementary Fig. 1: All GEPs in blood and liver.** Violin plots show GEP usage between cells from participants with paired blood (red) and liver (blue) cells. Statistical significance was determined using Wilcoxon rank-sum test (unpaired, nonparametric, two-sided) and for DGE supplemented with Benjamini-Hochberg correction for multiple testing; **p* ≤ 0.05, ** *p* ≤ 0.01, *** *p* ≤ 0.001, non-significant *p*-value is given if *p* > 0.05.

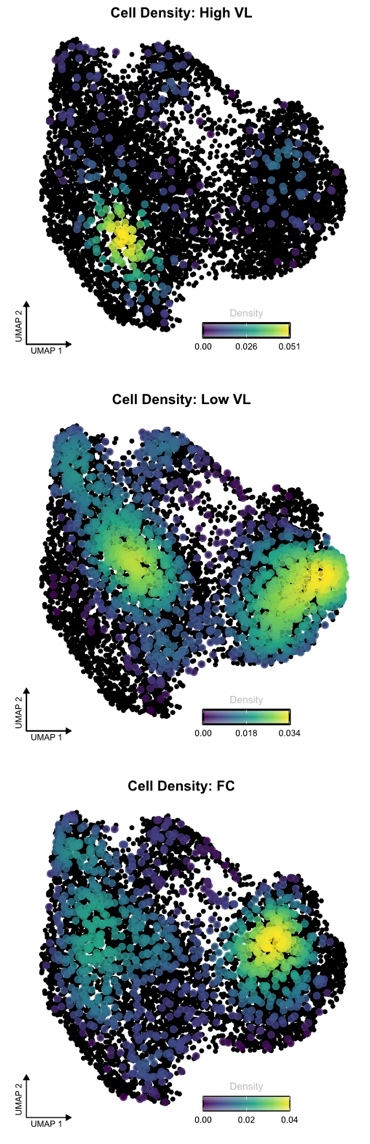

**Supplementary Fig. 2: Cells in CHB stages.** UMAP shows density of cells from the indicated cohorts of CHB and FC.

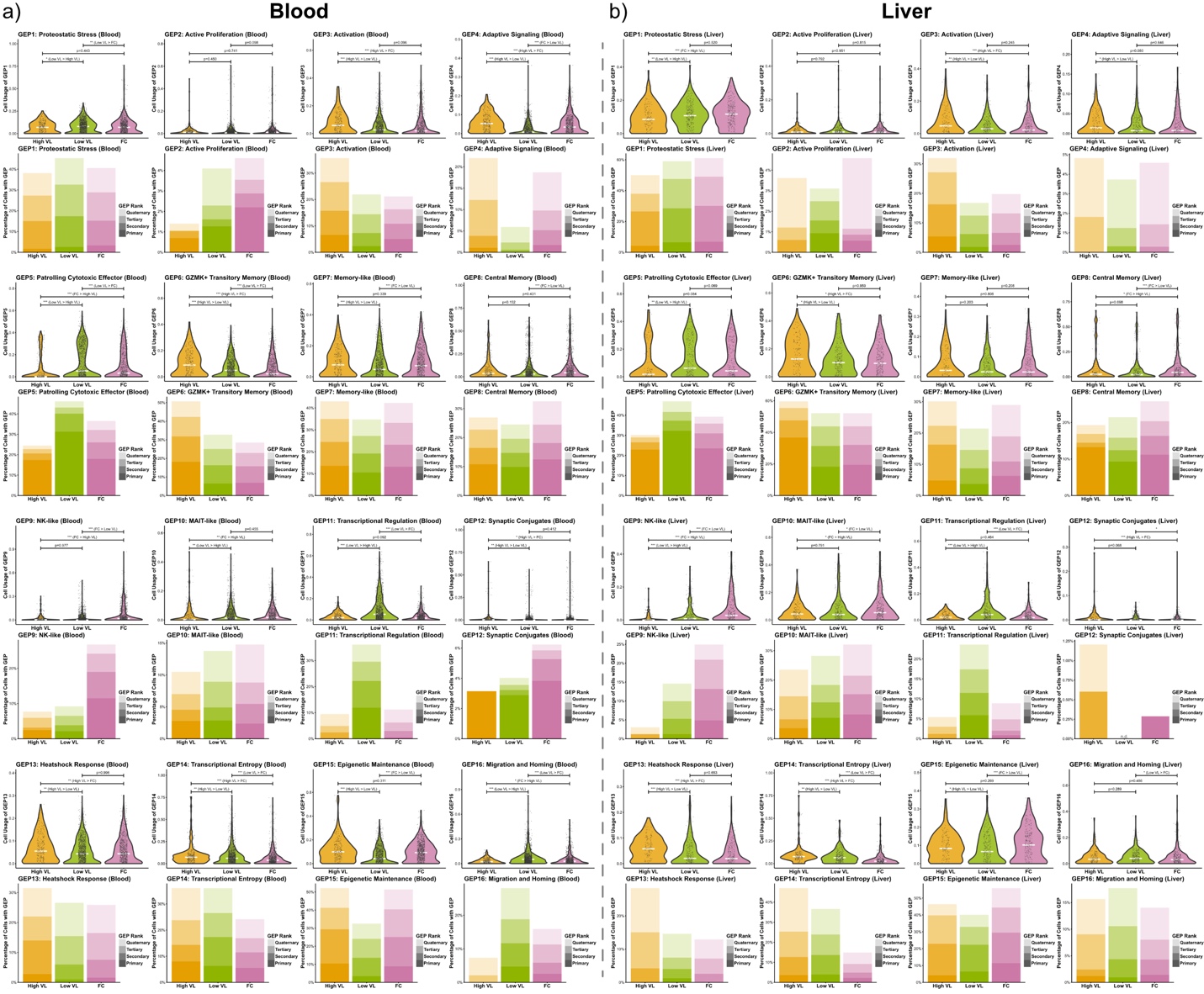

**Supplementary Fig. 3: All GEPs in CHB high VL, low VL and FC.** (**a**) Cells from blood, and (**b**) cells from liver are shown. Violin plots show GEP usage between cells from participants with CHB high VL (orange), low VL (green) and FC (purple). Hierarchical stacked bar chart shows abundance of cells with respective GEP as their primary highest usage, secondary, tertiary, or quaternary GEP. Statistical significance was determined using Wilcoxon rank-sum test (unpaired, nonparametric, two-sided) and for DGE supplemented with Benjamini-Hochberg correction for multiple testing; **p* ≤ 0.05, ** *p* ≤ 0.01, *** *p* ≤ 0.001, non-significant *p*-value is given if *p* > 0.05.

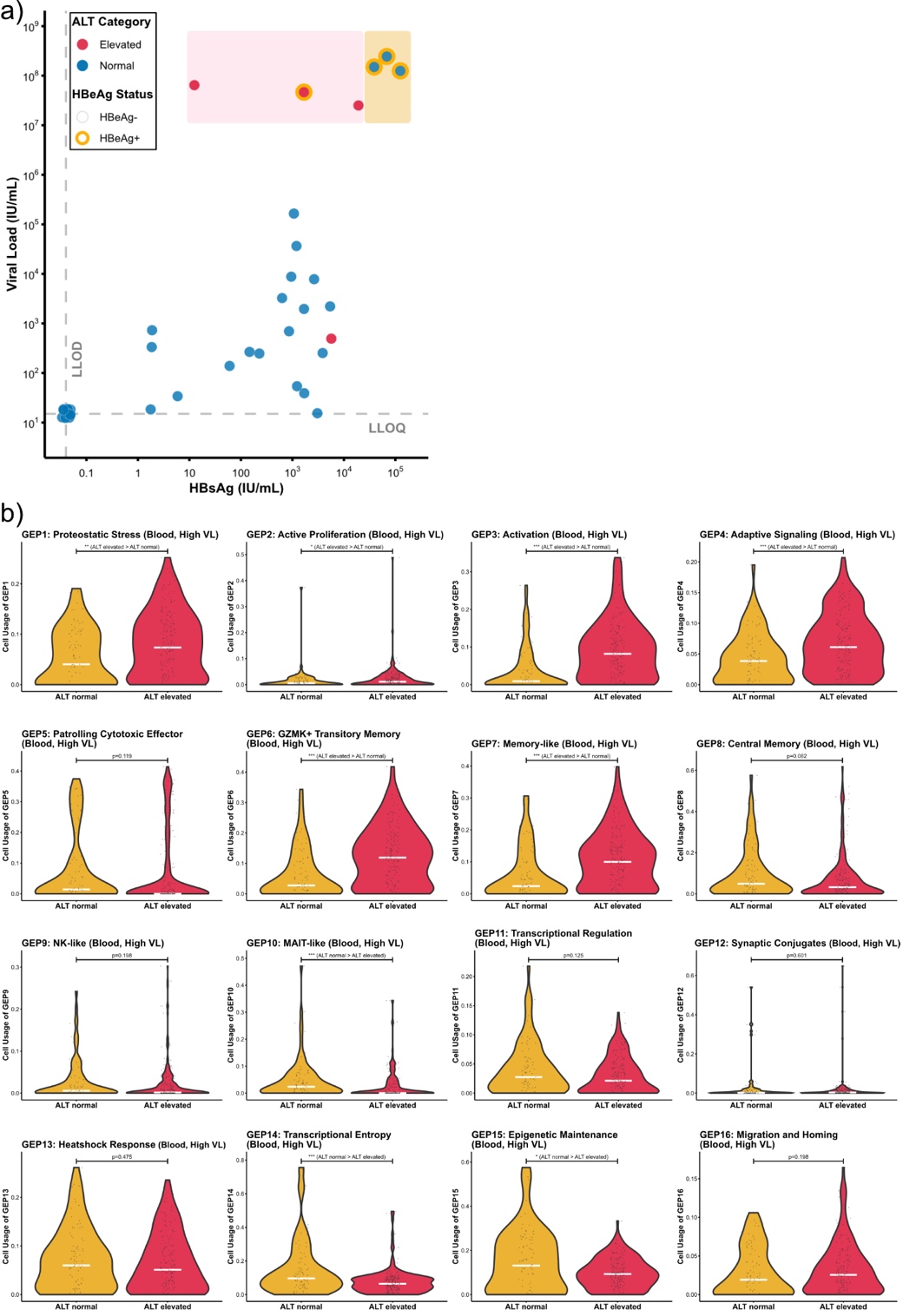

**Supplementary Fig. 4: Effect of ALT on T-cell GEP.** (**a**) Distribution of participants along their HBV viral load versus HBsAg. CHB participants with high viral load (≥ 1,000,000 IU/mL, normal ALT; yellow) are compared with elevated ALT and high viral load (≥ 1,000,000 IU/mL, elevated ALT; red). ALT category (elevated in red, normal in blue) and HBeAg status (orange circle is positive) are indicated. (n = 38 participants). (**b**) Violin plots show GEP usage between cells from participants with normal ALT (yellow) and elevated ALT (red). Statistical significance was determined using Wilcoxon rank-sum test (unpaired, nonparametric, two-sided) and for DGE supplemented with Benjamini-Hochberg correction for multiple testing; **p* ≤ 0.05, ** *p* ≤ 0.01, *** *p* ≤ 0.001, non-significant *p*-value is given if *p* > 0.05.

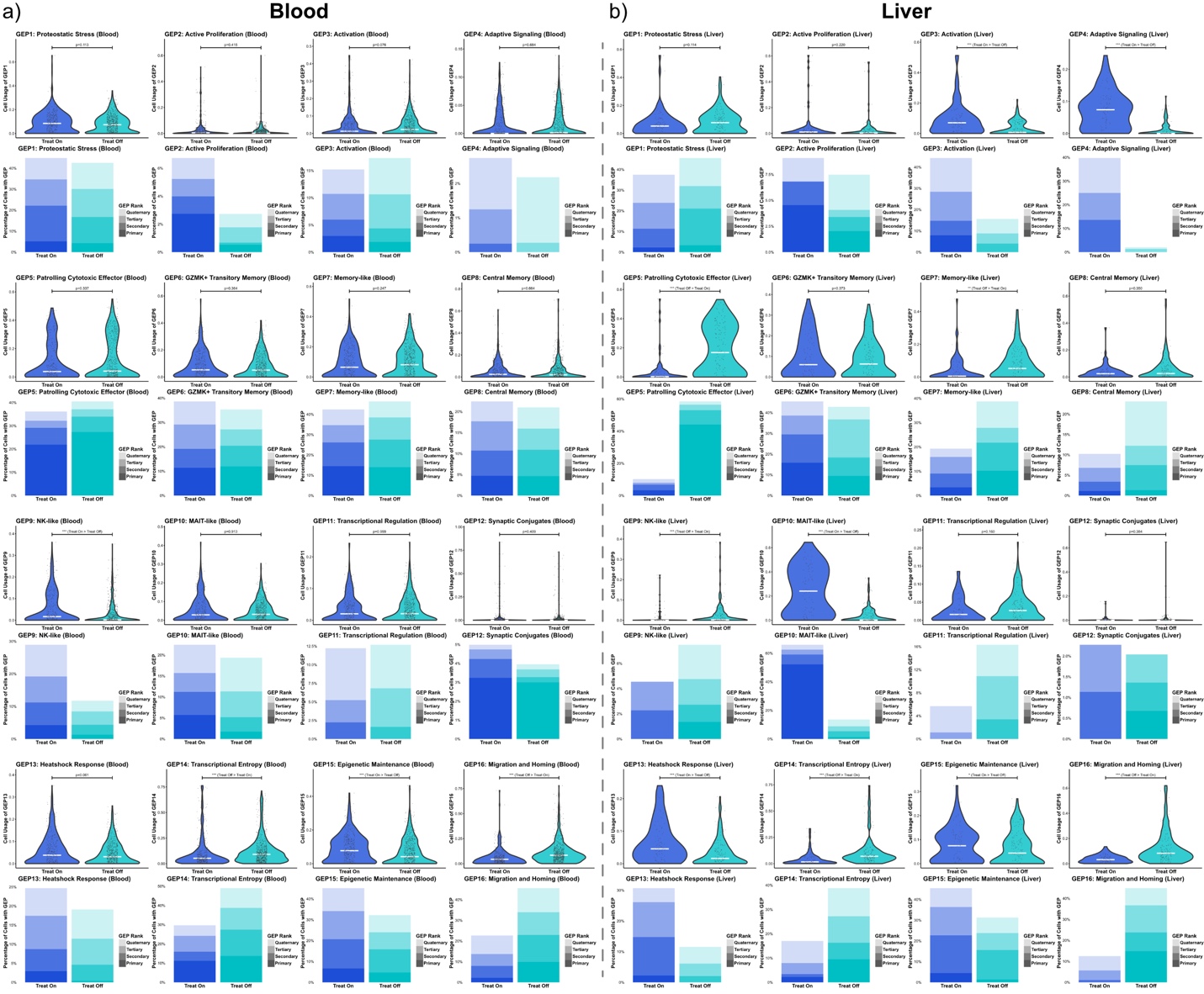

**Supplementary Fig. 5: All GEPs in Treated CHB.** (**a**) Cells from blood, and (**b**) cells from liver are shown. Violin plots show GEP usage between cells from participants with treated CHB (blue), and treatment interruption (cyan). Hierarchical stacked bar chart shows abundance of cells with respective GEP as their primary highest usage, secondary, tertiary, or quaternary GEP. Statistical significance was determined using Wilcoxon rank-sum test (unpaired, nonparametric, two-sided) and for DGE supplemented with Benjamini-Hochberg correction for multiple testing; **p* ≤ 0.05, ** *p* ≤ 0.01, *** *p* ≤ 0.001, non-significant *p*-value is given if *p* > 0.05.

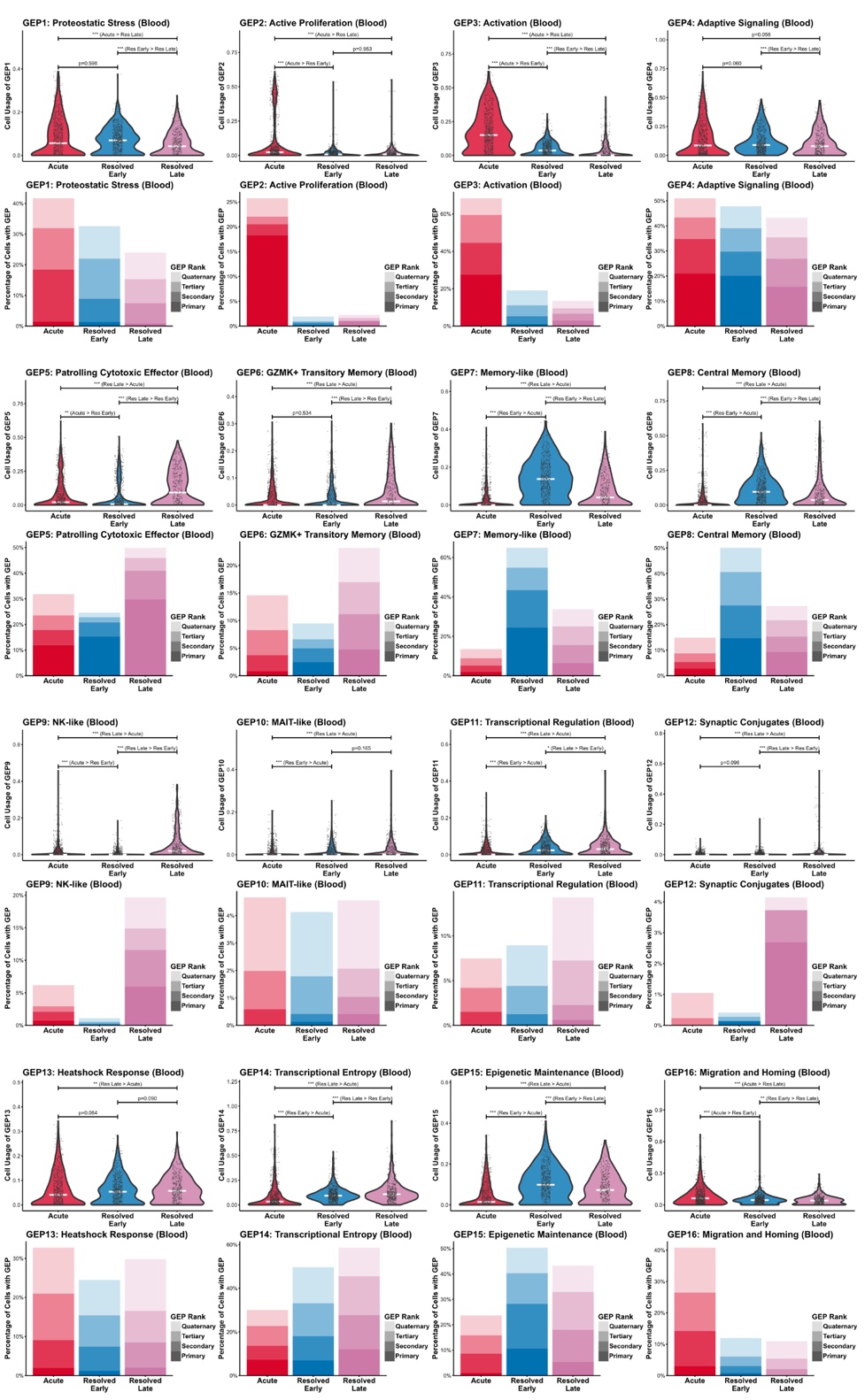

**Supplementary Fig. 6: All GEPs in Acute HBV infection.** Violin plots show GEP usage between blood-derived cells from participants with acute (red), resolved acute HBV infection < 1 year of resolution (blue), and resolved acute HBV infection ≥ 1 year of resolution (purple). Hierarchical stacked bar chart shows abundance of cells with respective GEP as their primary highest usage, secondary, tertiary, or quaternary GEP. Statistical significance was determined using Wilcoxon rank-sum test (unpaired, nonparametric, two-sided) and for DGE supplemented with Benjamini-Hochberg correction for multiple testing; **p* ≤ 0.05, ** *p* ≤ 0.01, *** *p* ≤ 0.001, non-significant *p*-value is given if *p* > 0.05.

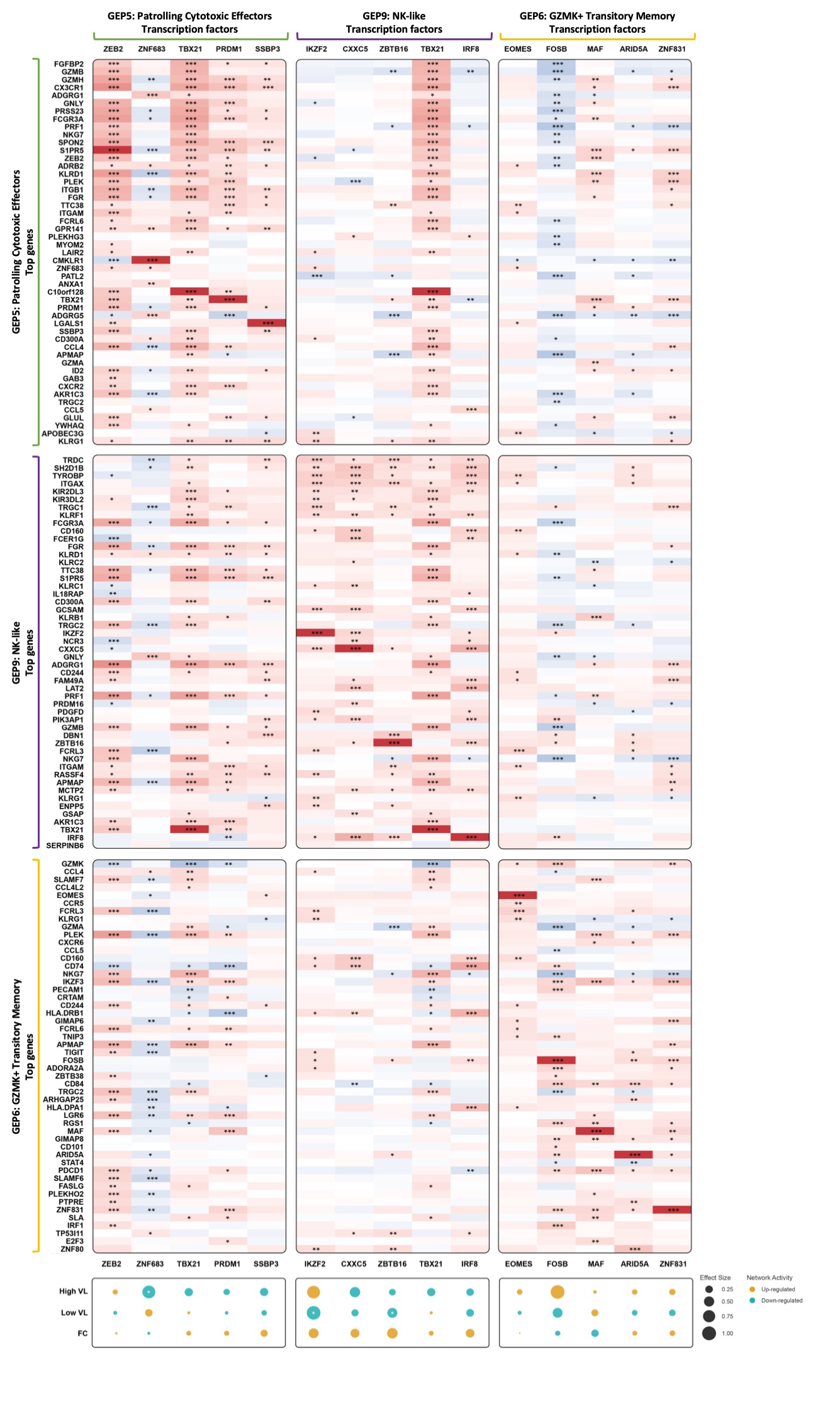

**Supplementary Fig. 7: Transcription factor enrichment of different cytotoxic T-cell programs.** Heatmap with correlation between top5 transcription factors (x-axis) of three effector GEPs *Patrolling Cytotoxic Effector* (left), *NK-like* (middle), and *GZMK+ Transitory Memory* (right) with top50 genes (y-axis). Dotplot shows network activity according to Gene Regulatory Network for the indicated TF within the GEP for HBV-specific CD8 T cells from CHB participants with High VL, Low VL, and FC. Effect size is indicated by the size of the dot. The effect indicates either an up-regulation (yellow), or a down-regulation (teal). Correlation was determined using Spearman’s rank correlation. Statistical significance was determined using Wilcoxon rank-sum test (unpaired, nonparametric, two-sided) with Benjamini-Hochberg correction for multiple testing; **p* ≤ 0.05, ** *p* ≤ 0.01, *** *p* ≤ 0.001, non-significant *p*-value is given if *p* > 0.05.

Legends for Supplementary Tables:

**Supplementary Table 1:** Overview of multimers used in this study.

**Supplementary Table 2:** The clinical characteristics and the information on immunological assays of the individuals in the chronic hepatitis B cohort.

**Supplementary Table 3:** Top200 genes of all 16 GEP.

**Supplementary Table 4:** DGE of the different comparisons (as depicted in volcano plots) throughout the manuscript are shown with fold change, *p*-value, and adjusted *p*-value. Genes considered to be significant are indicated on which comparator they are upregulated, otherwise they are listed as non-significant.

**Supplementary Table 5:** The clinical characteristics of the individuals in the longitudinal treatment-interruption hepatitis B cohort.

**Supplementary Table 6:** The clinical characteristics of the individuals in the acute hepatitis B and acute-controlled cohort.

**Supplementary Table 7:** The antibody-fluorochrome combinations used in this study.
